# Prefrontal somatostatin interneurons encode fear memory

**DOI:** 10.1101/696377

**Authors:** Kirstie A. Cummings, Roger L. Clem

## Abstract

Theories stipulate that memories are encoded within networks of cortical projection neurons (PNs). Conversely, GABAergic interneurons (INs) are thought to function primarily to inhibit PNs and thereby impose network gain control, an important but purely modulatory role. However, we found that associative fear learning potentiates synaptic transmission and cue-specific activity of medial prefrontal cortex (mPFC) somatostatin interneurons (SST-INs), and that activation of these cells controls both memory encoding and expression. Furthermore, the synaptic organization of SST- and parvalbumin (PV)-INs provides a potential circuit basis for SST-IN-evoked disinhibition of mPFC output neurons and recruitment of remote brain regions associated with defensive behavior. These data suggest that rather than constrain mnemonic processing, potentiation of SST-IN activity represents an important causal mechanism for conditioned fear.

## Main

Associative memory is a critical function of cortical brain networks, which are primarily populated by excitatory PNs and inhibitory INs. The most abundant of these cell types, PNs are a key substrate for interregional brain signaling that is critical for memory expression ^1,2^. Accordingly, retrieval cues activate subsets of PNs that are hypothesized to encode stimulus associations through persistent changes in excitatory synapse strength and density ^3^. In contrast, GABAergic INs are generally thought to inhibit PNs ^4–8^, which has been suggested to optimize the dynamic range of PN firing to indirectly modulate the strength and specificity of learning. While several studies credit INs with such “supporting roles”, it remains unclear whether they can directly mediate the encoding of cue associations through their own functional plasticity.

Fear conditioning is a powerful model of such learning in which an animal acquires survival-based defensive reactions to a conditioned stimulus (CS) that predicts imminent threat. The expression of fear memory in rodents requires neural activity in the prelimbic subregion of mPFC ^9^, where both PNs and INs sampled by extracellular recordings exhibit CS-evoked changes in firing rate after conditioning ^4^. However, it is unknown whether learning induces synaptic plasticity in prelimbic circuits and, if so, whether IN activity is modulated by these changes in conjunction with memory encoding. Here we address these questions in mice with a combination of synaptic electrophysiology, calcium imaging, optogenetic manipulation and brain activity mapping of prelimbic interneurons and associated circuitry. We demonstrate that SST-INs exhibit properties indicative of a memory storage substrate, including **1.** learning-dependent potentiation of synaptic transmission, **2.** cue-specific activation during memory retrieval, and **3.** bidirectional modulation of memory expression. Moreover, prelimbic SST-INs exert potent disinhibitory control over a fear-related brain network, suggesting a fundamental role for these cells in orchestrating conditioned fear responses.

## Results

### Cued fear learning potentiates SST-IN excitatory input

While experience-dependent plasticity is considered to be the most likely mechanism for cortical information storage ^2,3^, the extent to which learning is associated with plasticity of cortical inhibitory circuits remains poorly understood. To determine whether fear learning alters the synaptic properties of prefrontal INs, we therefore obtained *ex vivo* electrophysiological recordings from parvalbumin (PV)- and SST-expressing cells, which together comprise the majority of cortical GABAergic interneurons ^10^. In order to identify these cell types in acute brain slices we generated IN-specific expression of tdTomato by crossing the Ai9 reporter line to PV- or SST-Cre driver mice, which exhibit highly selective recombination in prelimbic cortex (Fig. 1a)^11^. We then subjected these animals to behavioral training entailing either paired or unpaired presentations of an auditory CS and footshock (unconditioned stimulus, US; Fig. 1b; Supplementary Fig. 1a, f). Because only paired animals acquire CS-evoked defensive freezing, unpaired mice served as a control for non-associative effects of stimulus exposure ^12,13^. At 24 hours after training, spontaneous excitatory (sEPSCs) and inhibitory postsynaptic currents (sIPSCs) were recorded and analyzed as a proxy for potential synaptic plasticity in prelimbic INs (Fig. 1b-c; Supplementary Fig. 1). Mice that received CS-US pairing displayed higher sEPSC frequency in SST-but not PV-INs residing in layer 2/3 (L2/3), compared to naïve and unpaired controls (Fig. 1c). No other differences in sEPSC or sIPSC properties were associated specifically with CS-US pairing (Supplementary Fig. 1). Because sEPSC frequency can reflect differences in presynaptic efficacy, we next measured the response of layer 2/3 SST-INs to local paired-pulse stimulation, a well-established assay for neurotransmitter release probability (Fig. 1d) ^14^. Consistent with increased glutamate release onto SST-INs after fear conditioning, evoked EPSCs exhibited a higher paired-pulse ratio in animals that received CS-US pairing but not unpaired training. These results confirm that cued fear learning is associated with potentiation of excitatory synapses onto SST-INs.

**Fig. 1.**
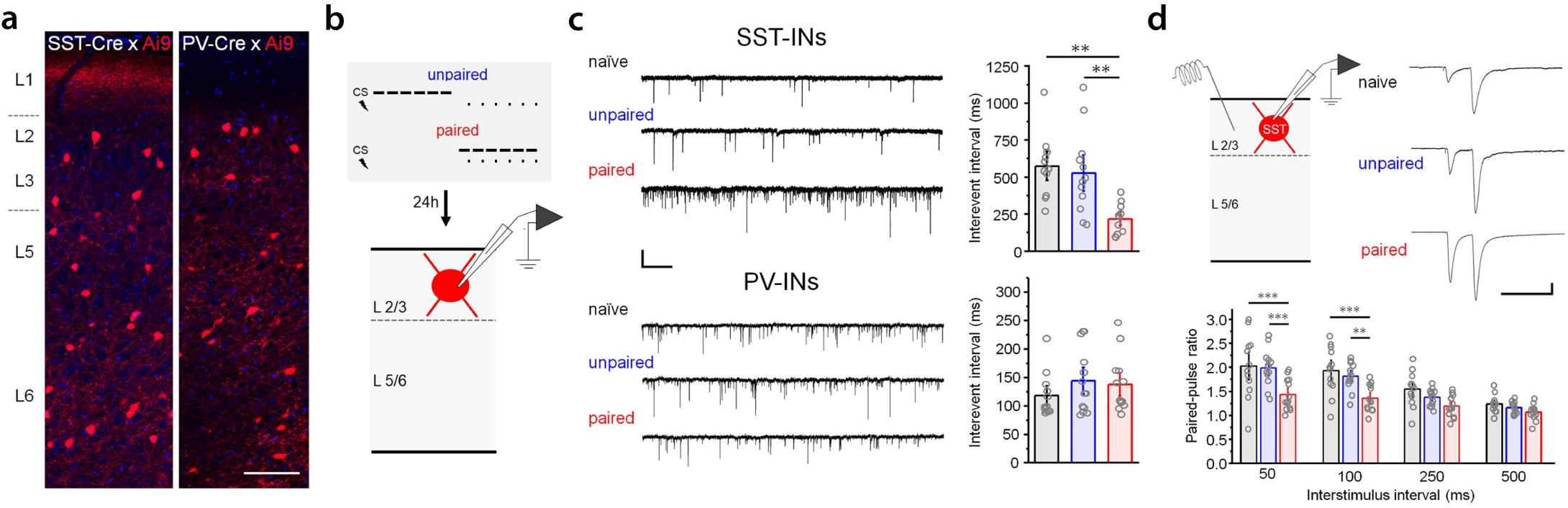
Potentiation of synaptic transmission in prefrontal SST-INs after cued fear learning. **a,** Prelimbic (PL) SST-INs and PV-INs were identified by tdTomato expression in SST-IRES-Cre/ Ai9 or PV-IRES-Cre/ Ai9 mice. Scale = 200 μm. **b,** Subjects were either cage-experienced (naïve), or trained using paired or unpaired presentations of CS (2 KHz, 80 dB, 20 s) and US (1 mA, 2-s), followed 24 hrs later by whole-cell recording in acute brain slices. **c,** Recordings of sEPSCs were obtained from L2/3 SST-INs (upper) and PV-INs (lower). Example raw traces and mean inter-event intervals are depicted for experimental and control groups. SST-INs: F_2,31_ = 9.03, p = 8.11 × 10^−4^, 1-way ANOVA; naïve, n = 11 cells (3 mice); unpaired n = 12 cells (3 mice); paired n = 11 cells (4 mice). PV-INs: F_2,45_ = 0.52, p = 0.60, 1-way ANOVA; naïve, n = 13 cells (3 mice); unpaired, n = 13 cells (3 mice); paired, n = 13 cells (4 mice). Scale = 20 pA × 1s. **d,** EPSC recordings in L2/3 SST-INs during paired-pulse stimulation. Paired pulse ratio: F_6,66_ = 3.46, p = 0.0049, interaction between ratio and training, 2-way repeated measures ANOVA; naïve, n = 13 cells (3 mice); unpaired n = 13 cells (3 mice); paired, n = 13 cells (3 mice). Scale = 100 pA x 100 ms. * p < 0.05, ** p < 0.01, *** p < 0.001 Tukey’s post-hoc test. Bar graphs depict mean ± SE.

### SST-INs signal the conditioned stimulus after learning

Given the observed learning-related potentiation of their excitatory input (Fig. 1), we hypothesized that SST-INs in conditioned mice might exhibit increased CS-related activity and thereby participate in memory processing. To test this possibility, we utilized fiber photometry to monitor *in vivo* activity of SST-INs after conditional viral expression of GCamp6f (AAV-DIO-GCamp6f-eYFP) in SST-Cre mice (Fig. 2; Supplementary Fig. 2). Mice were unilaterally implanted with an optic fiber (400 μm core diameter) directed at the prelimbic cortex and imaged under freely-behaving conditions during CS exposure before, during, and after auditory fear conditioning (Fig. 2b-c). During CS-US pairing, we observed an increase in CS-related calcium signals during trials 5-6 compared to the initial conditioning trial (Fig. 2d), which mirrored a trial-dependent increase in CS-evoked freezing (Fig. 2b). One day after conditioning, CS-related calcium signals during re-exposure to the CS were markedly increased compared to a pre-conditioning test (Fig. 2e). Because CS presentation leads to defensive freezing, the recruitment of SST-INs during memory retrieval could be a consequence of fear expression rather than CS modulation per se. We therefore additionally analyzed calcium signals during inter-trial freezing bouts, during which fear-related SST-IN activity can be dissociated from CS processing. Unlike CS trials, transitions from movement to freezing during the inter-trial intervals were associated with negligible fluorescence changes (Fig. 2e).

**Fig. 2.**
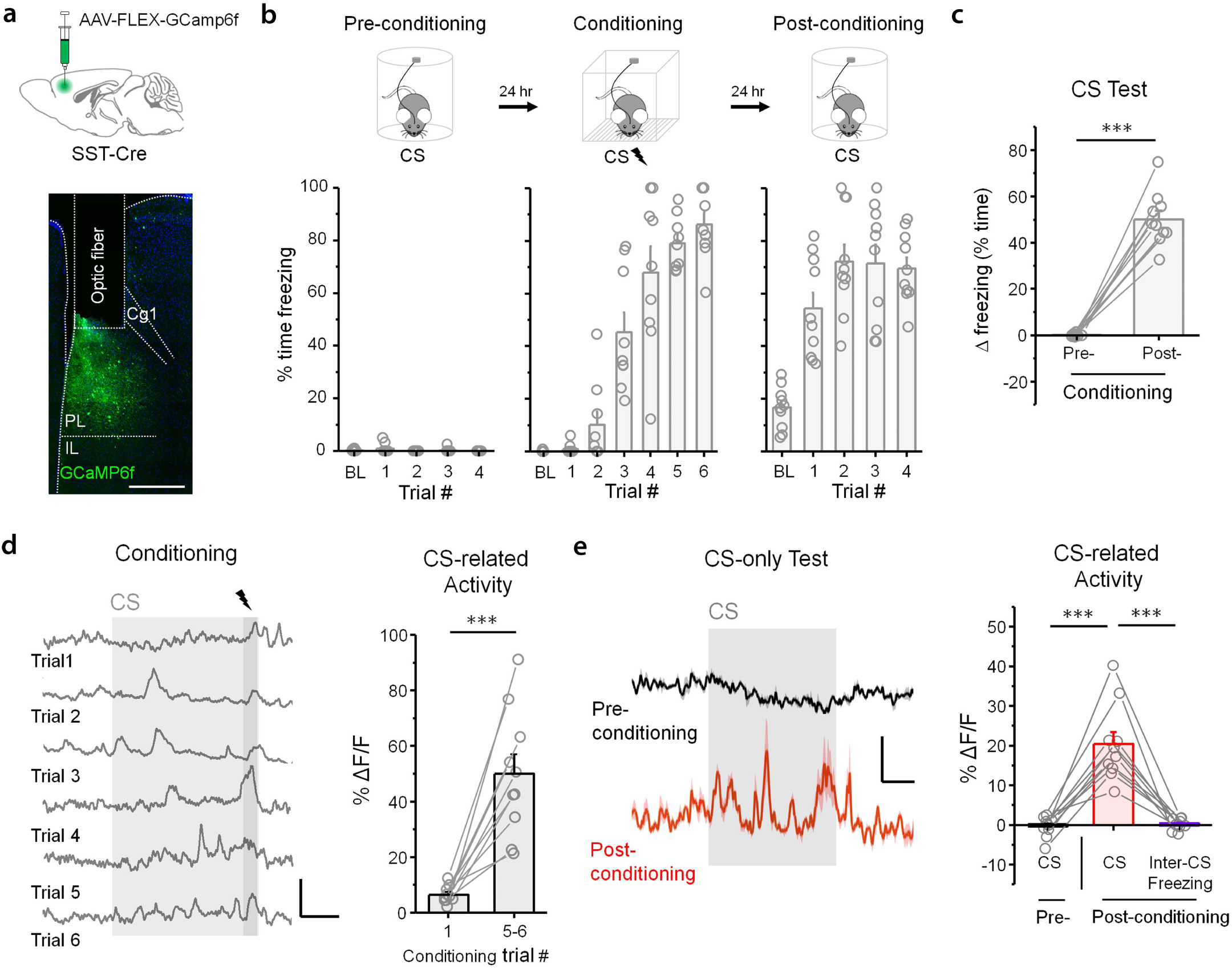
CS-related SST-IN activity increases in tandem with memory acquisition. **a**, For Ca^2+^-based imaging of SST-IN activity, SST-IRES-Cre mice (n = 10) received injections of conditional vector encoding GCamp6f and were implanted with a single optic fiber (400 μm core diameter) directed at PL. Scale = 500 μm. **b**, After surgical recovery, CS-related freezing and Ca^2+^-dependent fluorescence signal were monitored before, during and after CS-US pairing, which entailed 6 co-terminating trials of CS (2 KHz, 20 s, 80 dB) and US (0.7 mA foot shock, 2 s). Pre- and post-conditioning CS tests consisted of 4 CS trials in a context distinct from the training arena. **c**, Mean change in freezing induced by CS presentation during pre- and post-conditioning tests. Effect of training: t_9_ = 13.7. p = 2.51 × 10^−7^, paired t-test. **d**, CS-related fluorescence signal during CS-US pairing in a representative subject, and comparison of mean change in peak fluorescence (%ΔF/F) for conditioning trials 1 and trials 5-6. Effect of trial: t_9_ = 5.95, p = 2.15 × 10^−4^, paired t-test. Scale = 60% ΔF/F x 5 s. **e**, Mean CS-related fluorescence signal during pre- and post-conditioning CS tests in a representative subject. Right, comparison of percent ΔF/F between pre- and post-conditioning CS presentations, as well as freezing-related epochs that occurred independent of the CS during the inter-trial intervals of the post-conditioning test. CS-related activity: F_2,18_ = 42.82, p = 1.44 × 10^−7^, 1-way repeated measures ANOVA. Traces represent the average of 4 CS trials. Scale = 30% ΔF/F x 5 s. *** p < 0.001 by paired t-test (**c, d**) or Tukey’s post-hoc test (**e**). Bar graphs depict mean ± SE.

To further test whether fearful states modulate SST-IN activity independent of the CS, we next performed SST-IN-specific fiber photometry in mice that were subjected to unpaired CS-US training (Supplementary Fig. 3a-c). Importantly, although unpaired training results in context conditioned fear, it does not induce synaptic plasticity in SST-INs (Fig. 1c-d). During unpaired conditioning, US trials were associated with large calcium signals, confirming that changes in SST-IN activity could be readily detected (Supplementary Fig. 3d, f). However, regardless of whether imaging was conducted in a novel context (context B) or the original training arena (context A), no changes in calcium signals were associated with spontaneous freezing bouts (Supplementary Fig. 3e-f). Moreover, there were no overall differences in the frequency of calcium transients (peaks) during exposure to contexts A and B, relative to a preconditioning baseline in context A (Supplementary Fig. 3g), despite robust differences in freezing between these tests (Supplementary Fig. 3b). These data indicate that SST-INs do not generally signal a high-fear state, but are instead specifically activated by the threat-associated cue.

**Fig. 3.**
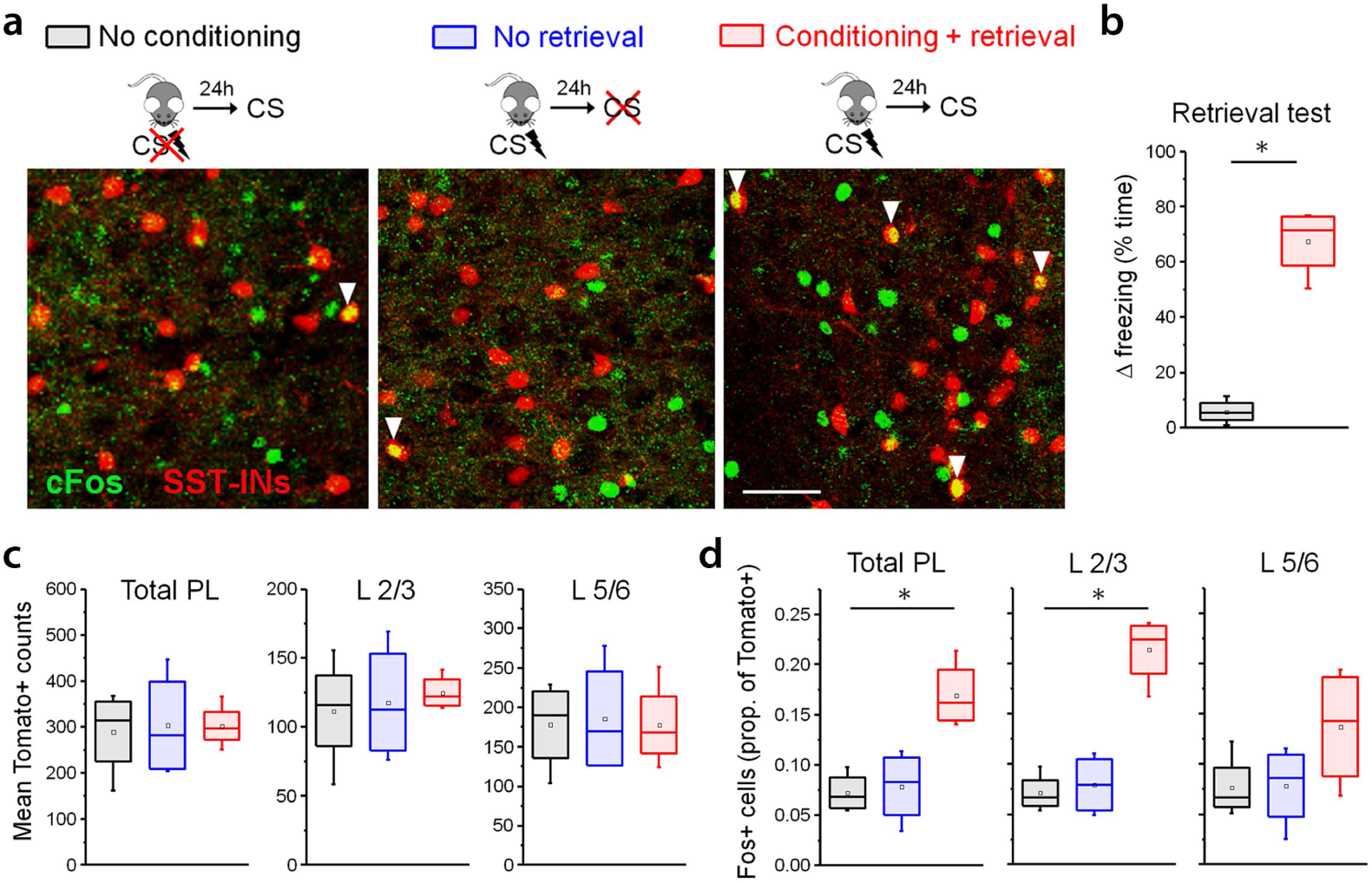
Fear learning increases cue-evoked cFos expression in SST-INs. **a,** cFos labeling in SST-INs following CS exposure (4 trials, 20 s duration) at 24 hrs after fear conditioning (n = 4 mice), as compared to control conditions in which conditioning (n = 4 mice) or CS exposure (n = 4 mice) were omitted. Arrowheads denote SST-INs co-labeled for both tdTomato (red) and cFos (green). **b,** CS-evoked freezing during the retrieval test for experimental animals and no conditioning controls. Tone test: *U* = 0, p = 0.030, Mann-Whitney *U* test. **c,** Mean number of Tomato+ SST-INs that were counted per brain section. **d,** Proportion of Tomato^+^ SST-INs that were co-labeled for cFos in each group. Total prelimbic (PL) counts: *χ*2 = 7.54 (2), p = 0.011, Kruskal-Wallis ANOVA. L2/3 counts: *χ*2 = 7.42 (2), p = 0.013, Kruskal-Wallis ANOVA. L5/6: *χ*2 = 2.35 (2), p = 0.35, Kruskal-Wallis ANOVA. * p < 0.05 Mann-Whitney *U* test **(b)**. * p < 0.05 Dunn’s post-hoc test in **(d)**. Box plots depict median (center line), mean (black box), quartiles, and 10-90% range (whiskers).

To independently confirm the activation of SST-INs in response to memory retrieval, we performed immunohistochemical labeling for the activity reporter cFos (Fig. 3a; Supplementary Fig. 2), which is considered to be a marker of neurons strongly activated by mnemonic cues ^3^. To elicit memory retrieval in SST-Cre/ Ai9 reporter mice, 4 CS trials were presented at 24 hours after fear conditioning in a context distinct from the training arena, and as control conditions we examined mice in which either conditioning or memory retrieval were omitted. Retrieval CS trials triggered a robust increase in freezing only in mice that had been previously conditioned (Fig. 3b; Supplementary Fig. 4). Following behavioral testing, substantially more SST-INs exhibited cFos immunoreactivity in these animals compared to those in either control group (Fig. 3c-d). Increased SST-IN activation in conditioned mice could also be observed specifically in layer 2/3, consistent with a causal role for lamina-specific SST-IN plasticity (Fig. 1). In contrast to these results, PV-INs did not exhibit a detectable increase in cFos immunoreactivity under the same conditions in PV-Cre/ Ai9 mice (Supplementary Fig. 5).

**Fig. 4.**
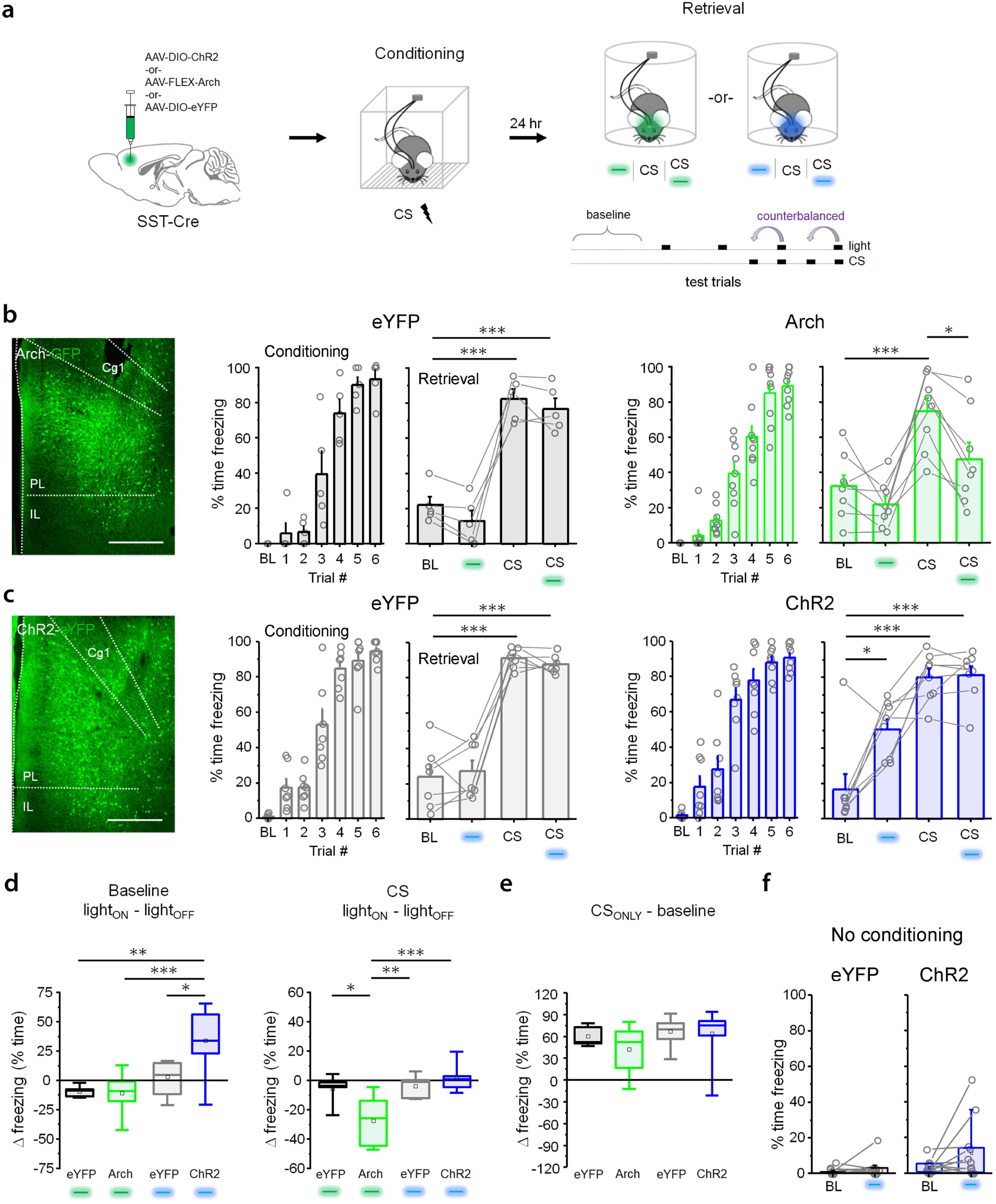
SST-IN activity is necessary for memory expression and sufficient to induce freezing in conditioned mice. **a,** For *in vivo* activity manipulation, SST-IRES-Cre mice received injections of conditional Arch, ChR2, or eYFP control vectors and were implanted with optic ferrules directed at PL. After surgical recovery, all subjects underwent auditory fear conditioning in the absence of optic illumination. Freezing was quantified 24 hrs later during independent and simultaneous presentation of CS and photostimulation in a distinct context. **b,** Modulation of freezing by CS and light (532 nm, constant, 20 s epochs) in Arch (green) or eYFP control mice (black). Arch: F_3,21_ = 11.3, p = 1.29 × 10^−4^, 1-way repeated measures ANOVA; n = 8 mice. eYFP: F_3,12_ = 68.6, p = 8.03 × 10^−8^, repeated measures ANOVA; n = 5 mice. Scale = 500 μm. **c,** Modulation of freezing by CS and light (473 nm, 10 ms pulses at 20 Hz) in ChR2 (blue) or eYFP control mice (gray). ChR2: F_3,21_ = 19.3, p = 3.06 × 10^−6^, 1-way repeated measures ANOVA; n = 9 mice. eYFP: F_3,18_ = 87.7, p = 6.20 × 10^−11^, 1-way repeated measures ANOVA; n = 7 mice. Scale = 500 μm. **d,** Mean change in freezing induced by photostimulation during baseline and CS periods, as compared across all light/ opsin combinations used in (**b**) and (**c**). Photostimulation effect on baseline freezing (light_ON_ – light_OFF_): F_3,24_ = 8.88, p = 3.87 × 10^−4^, 1-way ANOVA. Photostimulation effect on CS-evoked freezing (light_ON_ – light_OFF_): F_3,24_ = 9.61, p = 2.37 × 10^−4^, 1-way ANOVA. **e,** Mean change in freezing induced by CS presentation, as compared across all experimental groups used in (**b**) and (**c**). CS-evoked change in freezing (CS_ONLY_ – baseline): F_3,24_ = 1.12, p = 0.36, 1-way ANOVA. **f,** Freezing behavior during light-only trials (473 nm, 10 ms pulses, 20 Hz) in naïve (unconditioned) mice expressing either ChR2 or eYFP control vector. ChR2: *W* = 17, p = 0.088, Wilcoxon signed ranks test; n = 12 mice. eYFP: *W* = 12, p = 0.46, Wilcoxon signed ranks test; n = 11 mice. BL = baseline period. CS = conditioned stimulus (2 kHz tone, 80 db, 20 s duration). Cg1 = cingulate area 1. IL = infralimbic cortex. Blue/ green boxes = laser illumination. * p < 0.05, ** p < 0.01, ** p < 0.001 Tukey’s post-hoc test (**b, c, d**). Bar graphs depict mean ± SE. Box plots depict median (center line), mean (black box), quartiles, and 10-90% range (whiskers).

**Fig. 5.**
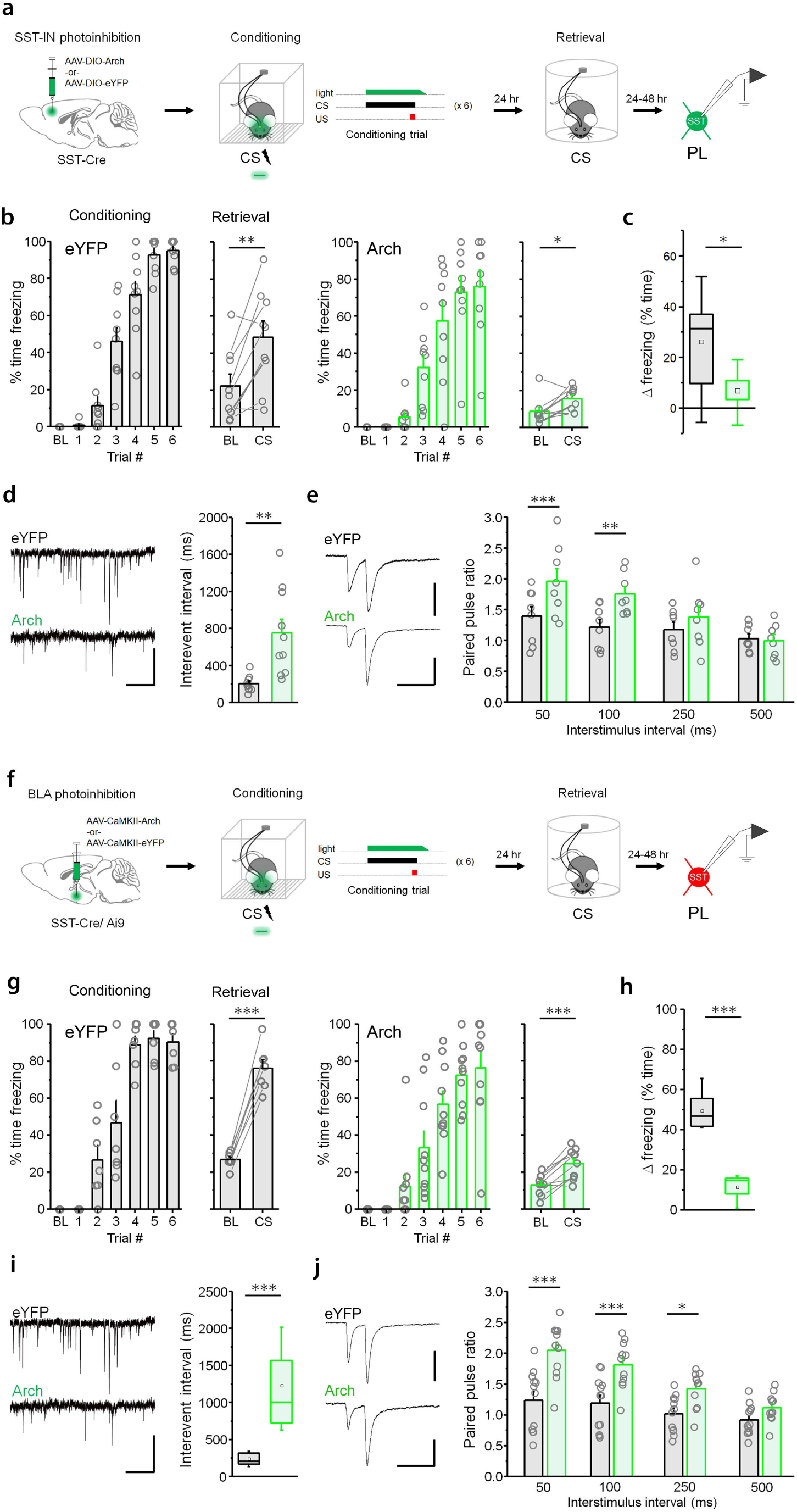
SST-IN activation and plasticity mediates memory formation. **a**, For *in vivo* manipulation of prelimbic SST-INs, SST-IRES-Cre mice received injections of conditional Arch or eYFP control vectors and were implanted with optic ferrules directed at PL. After surgical recovery, all subjects underwent auditory fear conditioning, during which light stimulation (532 nm, constant, 20 s epoch, ramp offset) coincided with each of 6 CS-US trials. Freezing was quantified 24 hrs later during presentation of 4 CS trials context distinct from the training arena. During the 24-48 hours subsequent to the retrieval test, *ex vivo* recordings were obtained from eYFP-positive neurons (SST-INs) located not more than 1 mm below the tip of the optic fiber in layer 2/3 of acute brain slices. **b**, Modulation of freezing by CS retrieval in Arch (green) or eYFP control mice (black). Arch retrieval: t_8_ = 2.71, p = 0.026, paired t-test; n = 9 mice. eYFP retrieval: t_8_ = 3.78, p = 0.0054, paired t-test; n = 9 mice. **c**, Mean change in freezing induced by CS presentation during memory retrieval for experimental groups in (**b**). Arch versus eYFP: t_16_ = 2.60, p = 0.019, unpaired t-test. **d**, Example raw traces and interevent intervals (IEIs) for sEPSCs in Arch versus eYFP mice. IEI: t_16_ = 2.60, p = 0.019, unpaired t-test; Arch, n = 10 cells (3 mice); eYFP, n = 10 cells (3 mice). Scale = 10 pA x 0.5 s. **e**, EPSC recordings in SST-INs during local paired-pulse electrical stimulation. Example traces collected at 50 ms interstimulus interval. Paired pulse ratio: F_3,21_ = 5.55, p = 0.0058, interaction between ratio and training, 2-way repeated measures ANOVA; Arch, n = 8 cells (3 mice); eYFP, n = 8 cells (3 mice). Scale = 40 pA x 100 ms. **f**, Experimental design for *in vivo* manipulation of BLA projection neurons was the same as in (**a**), with the exception that vector injections and light stimulation were delivered to the BLA, and *ex vivo* recordings were obtained from Tomato-positive SST-INs in the prelimbic cortex of SST-IRES-Cre/ Ai9 mice. **g**, Modulation of freezing by CS retrieval in Arch-(green) or eYFP control mice (black). Arch retrieval: t_9_ = 5.69, p = 2.98 × 10^−4^, paired t-test; n = 10 mice. eYFP retrieval: t_6_ = 14.74, 6.14 × 10^−6^, paired t-test; n = 7 mice. **h**, Mean change in freezing induced by CS presentation during memory retrieval for experimental groups in (**b**). Arch versus eYFP: *U* = 0, p = 1.03 × 10^−4^, Mann-Whitney *U* test. **i**, Example raw traces and interevent intervals (IEIs) for sEPSCs in Arch versus eYFP mice. IEI: *U* = 0, p = 9.97 × 10^−4^, Mann-Whitney *U* test; Arch, n = 14 cells (3 mice); eYFP, n = 13 cells (4 mice). Scale = 10 pA x 0.5 s. Scale = 10 pA x 0.5 s. **j**, EPSC recordings in SST-INs during local paired-pulse electrical stimulation. Example traces collected at 50 ms interstimulus interval. Paired pulse ratio: F_3,30_ = 11.18, p = 4.31 × 10^−5^, interaction between ratio and training, 2-way repeated measures ANOVA; Arch, n = 12 cells (3 mice); eYFP, n = 12 cells (3 mice). Scale = 50 pA x 100 ms. * p < 0.05, ** p < 0.01, *** p < 0.001 by paired t-test (**b, g**), unpaired t-test (**c, d**), Mann-Whitney *U* test (**h, i**), or Tukey’s post-hoc test (**j**). Bar graphs depict mean ± SE. Box plots depict median (center line), mean (black box), quartiles, and 10-90% range (whiskers).

### SST-IN activity controls memory expression

CS activation of SST-INs suggests that recruitment of these cells could be important for memory retrieval. On the other hand, previous studies suggest that SST-IN activity might also function to constrain fear expression through inhibition of cue-responsive PNs ^4–8^. To establish the behavioral impact of SST-IN recruitment, we utilized optogenetics to modulate SST-IN activity in conjunction with CS-evoked memory retrieval. We microinjected into prelimbic cortex viral vectors encoding archearhodopsin (Arch; AAV-FLEX-Arch3.0-GFP), channelrhodopsin-2 (ChR2; AAV-DIO-ChR2-eYFP), or an opsin-negative control vector (eYFP; AAV-DIO-eYFP). Importantly, optic illumination was sufficient to hyperpolarize or depolarize Arch- or ChR2-expressing SST-INs, respectively, and thereby reliably control the firing of these cells (Supplementary Fig. 6a-b). Following virus infusion, mice were implanted with optic fibers directed at the prelimbic cortex (Fig. 4a; Supplementary Fig. 6c-d). One week after surgery, these mice were subjected to CS-US pairing in the absence of photostimulation.

**Fig. 6.**
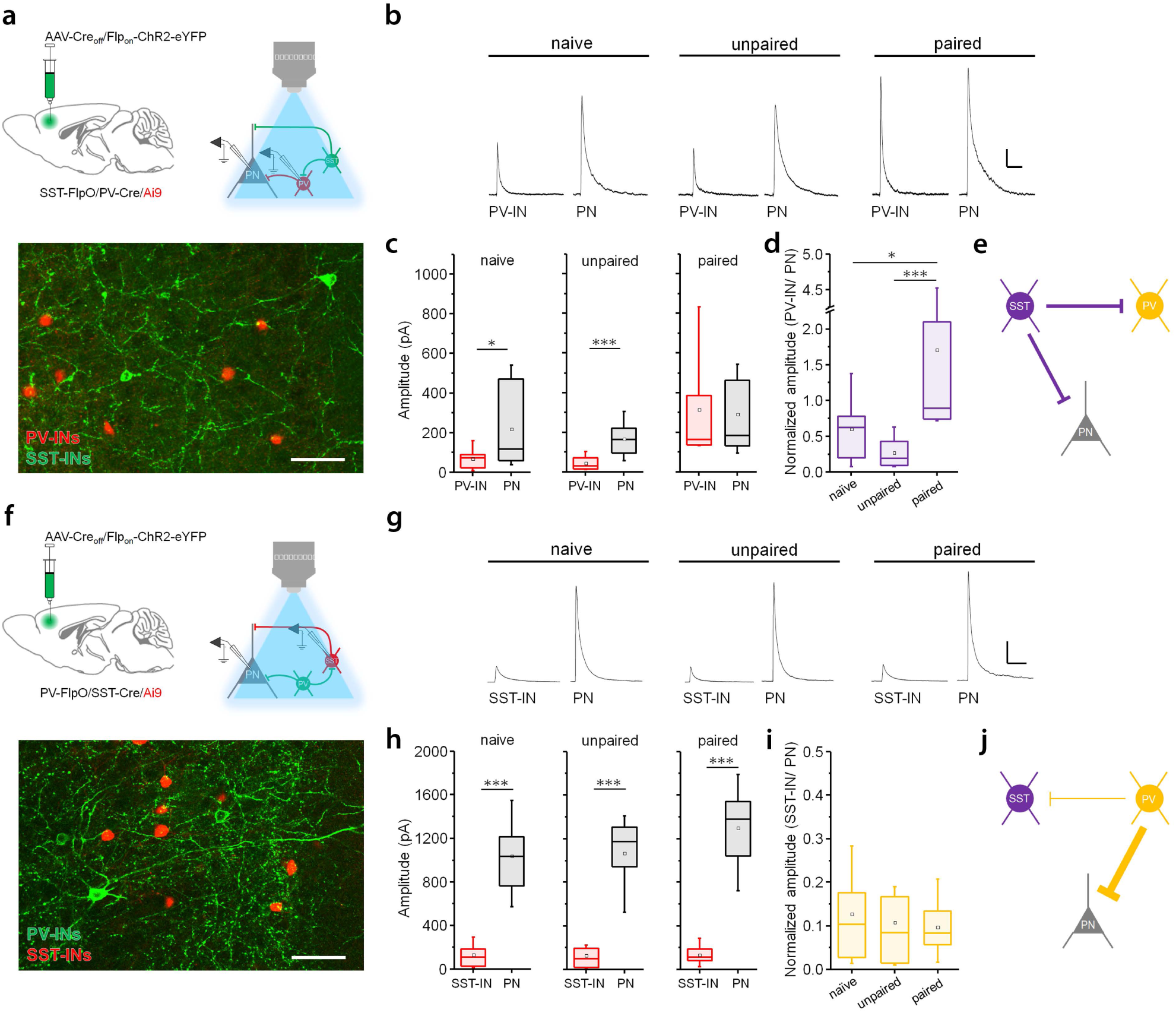
SST-INs elicit relatively potent inhibition of PV-INs while PV-INs preferentially inhibit PNs. **a,** To isolate monosynaptic responses to SST-IN photostimulation in PV-INs, we first infused a Flp-ON, Cre-OFF INTRSECT ChR2 vector into SST-IRES-FlpO/ PV-IRES-Cre/ Ai9 triple transgenic mice. At 24 hours after CS-US pairing, recordings were obtained from Tomato^+^ PV-INs as well as surrounding PNs. Scale = 50 μm. **b**, Example IPSC traces. Scale = 200 pA x 1 s. **c,** Amplitude of IPSCs resulting from SST-IN photoexcitation (460 nm, 1 ms pulse, 0.1 Hz) in slices from naïve mice (n = 4), as well as those that received unpaired (n = 3 mice) or paired training (n = 4 mice). Effect of cell type (naïve): *U* = 46.5, p = 0.017, Mann-Whitney *U* test; PV-IN, n = 14 cells; PN, n = 14 cells. Effect of cell type (unpaired): *U* = 17, p = 8.22 × 10^−4^, Mann-Whitney *U* test; PV-IN, n = 12 cells, PN, n = 12 cells. Effect of cell type (paired): *U* = 108.5, p = 0.88, Mann-Whitney *U* test; PV-IN, n = 15 cells; PN, n = 15 cells. **d**, Amplitude of IPSCs in PV-INs normalized to median values from PNs in the same slices. Effect of training: *χ*^2^ = 20.63 (2), p = 3.31 × 10^−5^, Kruskal-Wallis ANOVA. **e**, Relative strength of SST-IN transmission onto PV-INs versus PNs in conditioned mice. **f**, Interrogation of monosynaptic connections from PV-INs onto SST-INs and PNs. Flp-ON, Cre-OFF INTRSECT ChR2 vector into PV-IRES-FlpO/ SST-IRES-Cre/ Ai9 triple transgenic mice. **g**, Example IPSC traces. Scale = 200 pA x 1s. **h**, Amplitude of IPSCs resulting from PV-IN photoexcitation (460 nm, 1 ms pulse, 0.1 Hz) in slices from naïve mice (n = 3), as well as those that received unpaired (n = 3 mice) or paired training (n = 3 mice). Effect of cell type (naïve): *U* = 6, p = 1.15 × 10^−5^, Mann-Whitney *U* test; SST-IN, n = 14 cells; PN, n = 15 cells. Effect of cell type (unpaired): *U* = 3, p = 1.04 × 10^−5^, Mann-Whitney *U* test; PV-IN, n = 12 cells, PN, n = 11 cells. Effect of cell type (paired): *U* = 6, p = 1.15 × 10^−5^, Mann-Whitney *U* test; PV-IN, n = 12 cells; PN, n = 13 cells. **i**, Amplitude of IPSCs in PV-INs normalized to median values from PNs in the same slices. Effect of training: *χ*^2^ = 0.33 (2), p = 0.85, Kruskal-Wallis ANOVA. **j**, Relative strength of PV-IN transmission onto SST-INs versus PNs in conditioned mice. * p < 0.05, ** p < 0.01, *** p < 0.001 by Mann-Whitney *U* test (**c, h**) or Dunn’s post-hoc test (**d**). Box plots depict median (center line), mean (black box), quartiles, and 10-90% range (whiskers). During optic stimulation, 100% of postsynaptic cells that were sampled in these analyses exhibited synaptic responses.

At 24 hours after training, a memory retrieval test was conducted in which the independent and combined effects of light and CS were examined in a context distinct from the training arena. Compared to CS-only trials, a marked reduction in freezing was observed in Arch-expressing mice during light (532 nm, 20 s, constant) + CS trials (Fig. 4b,d), indicating that SST-IN activity is required for cued memory expression. No reduction in freezing was observed in Arch animals during light-only trials compared to the baseline period, suggesting that SST-IN activity is not required for generalized context fear. Conversely, in ChR2-expressing mice, photoexcitation of SST-INs (473 nm, 10 ms pulses, 20 Hz) was on its own sufficient to increase freezing over baseline levels and thereby mimic a conditioned response (Fig. 4c,d). Combined CS + light presentation in these animals did not increase freezing beyond that observed during interleaved CS-only trials, suggesting the possibility of a ceiling effect. Importantly, light-evoked freezing cannot be explained by a nonspecific motor deficit because photoexcitation of ChR2-expressing SST-INs did not alter standard metrics of locomotion in the open field test (Supplementary Fig. 7). Finally, no light effects were observed in eYFP control groups that were stimulated with the same parameters used in Arch (Fig. 4b,d) or ChR2 mice (Fig. 4c-d), and all groups displayed a CS-evoked increase in freezing of similar magnitude and therefore did not differ in memory strength (Fig. 4e).

**Fig. 7.**
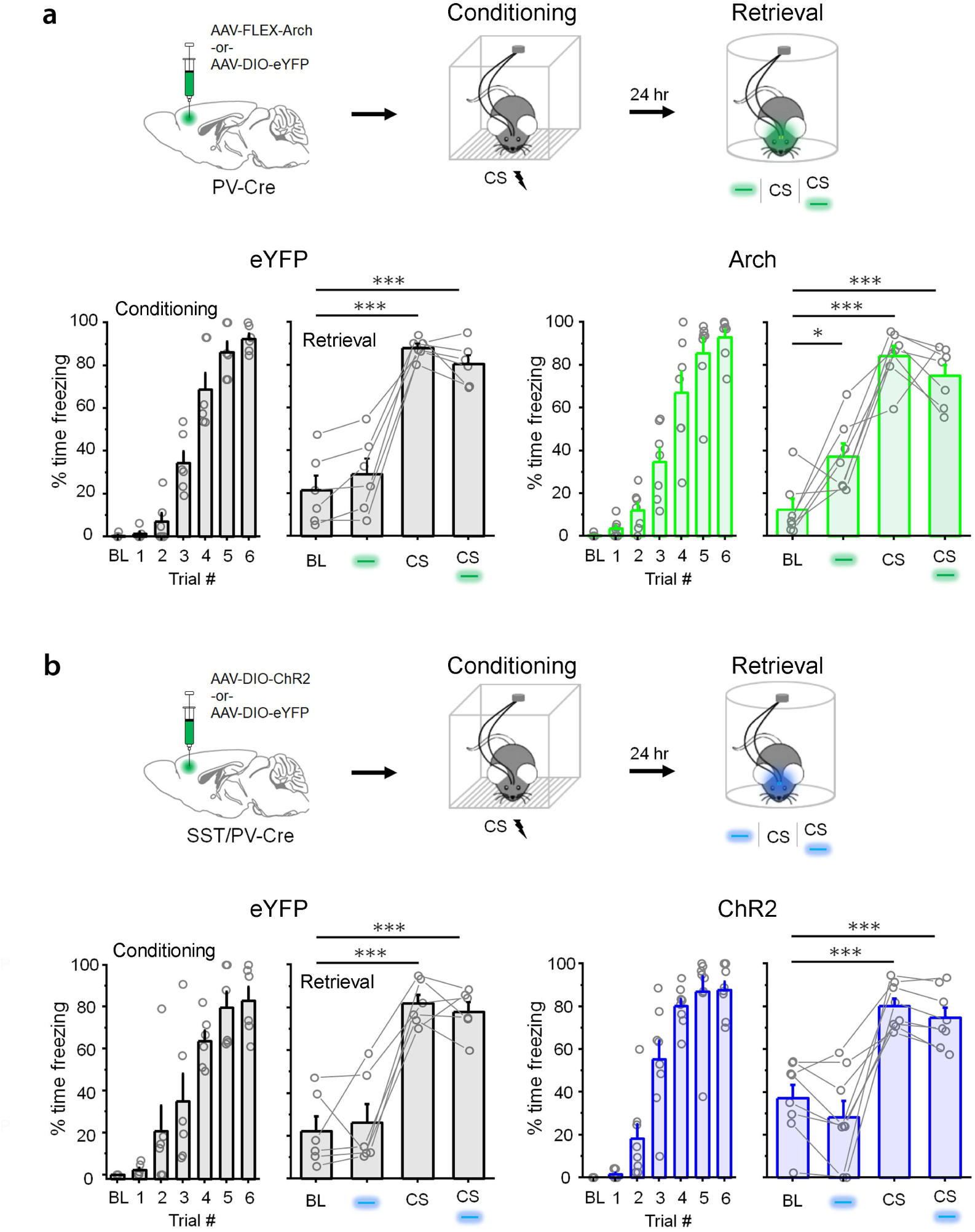
SST-IN-evoked freezing requires suppression of PV-INs. **a-b,** *In vivo* dissection of PV-IN contributions to stimulus-dependent freezing. **a,** Modulation of freezing by CS and light (532 nm, constant, 20 s) at 24 hrs after CS-US pairing in PV-IRES-Cre mice expressing conditional Arch (green) or eYFP control vectors (black). Arch: F_3,18_ = 33.2, p = 1.52 × 10^−7^, 1-way repeated measures ANOVA; n = 6 mice. eYFP: F_3,15_ = 50.1, p = 4.74 × 10^−8^, 1-way repeated measures ANOVA; n = 7 mice. **b,** Modulation of freezing by CS and light (473 nm, 10 ms pulses, 20 Hz) at 24 hrs after fear conditioning in SST-IRES-Cre/ PV-IRES-Cre double transgenic mice expressing conditional ChR2 (blue) or eYFP control vectors (black). ChR2: F_3,21_ = 23.3, p = 6.94 × 10^−7^, 1-way repeated measures ANOVA; n = 8 mice. eYFP: F_3,15_ = 41.3, p = 1.74 × 10^−7^, 1-way repeated measures ANOVA; n = 6 mice.* p < 0.05, *** p < 0.001 by Tukey’s post-hoc test. Bar graphs depict mean ± SE.

To establish whether fear-promoting properties are intrinsic to SST-INs or acquired through learning, we performed photoexcitation of SST-INs without prior fear conditioning. In contrast to fear conditioned mice (Fig. 4c), no consistent effect of photoexcitation was observed in naïve subjects (Fig. 4f).

### SST-IN activation and plasticity mediates memory acquisition

Given the critical role of SST-INs in memory expression, we next sought to utilize photoinhibition to determine whether SST-IN activity at the time of CS-US pairing is required for memory acquisition as well as associated plasticity of SST-INs. After prelimbic infusion of AAV-FLEX-Arch3.0-GFP or eYFP control vectors, SST-Cre mice were subjected to CS-US pairing, during which photoinhibition was timed to coincide with each of the 6 CS-US trials (Fig. 5a; Supplementary Fig. 8). On the following day, memory retrieval was examined in the absence of photoinhibition during presentation of 4 CS retrieval trials. Remarkably, while CS presentations triggered an increase in freezing in both Arch-expressing mice as well as eYFP controls, the magnitude of CS-evoked increase in freezing was dramatically lower in Arch-expressing mice (Fig. 5b-c). This effect cannot be attributed solely to Arch expression, because a retrieval deficit was not observed in Arch-expressing mice that were conditioned without photoinhibition (Fig. 4e). Following memory retrieval, acute brain slices were obtained to compare excitatory synaptic transmission in prelimbic SST-INs of Arch versus eYFP subjects, focusing on layer 2/3 cells located within 1 mm below the optic fiber track. Compared to eYFP controls, Arch-expressing SST-INs exhibited lower sEPSC frequency as well as higher paired-pulse ratios of evoked EPSCs (Fig. 5d-e).

**Fig. 8:**
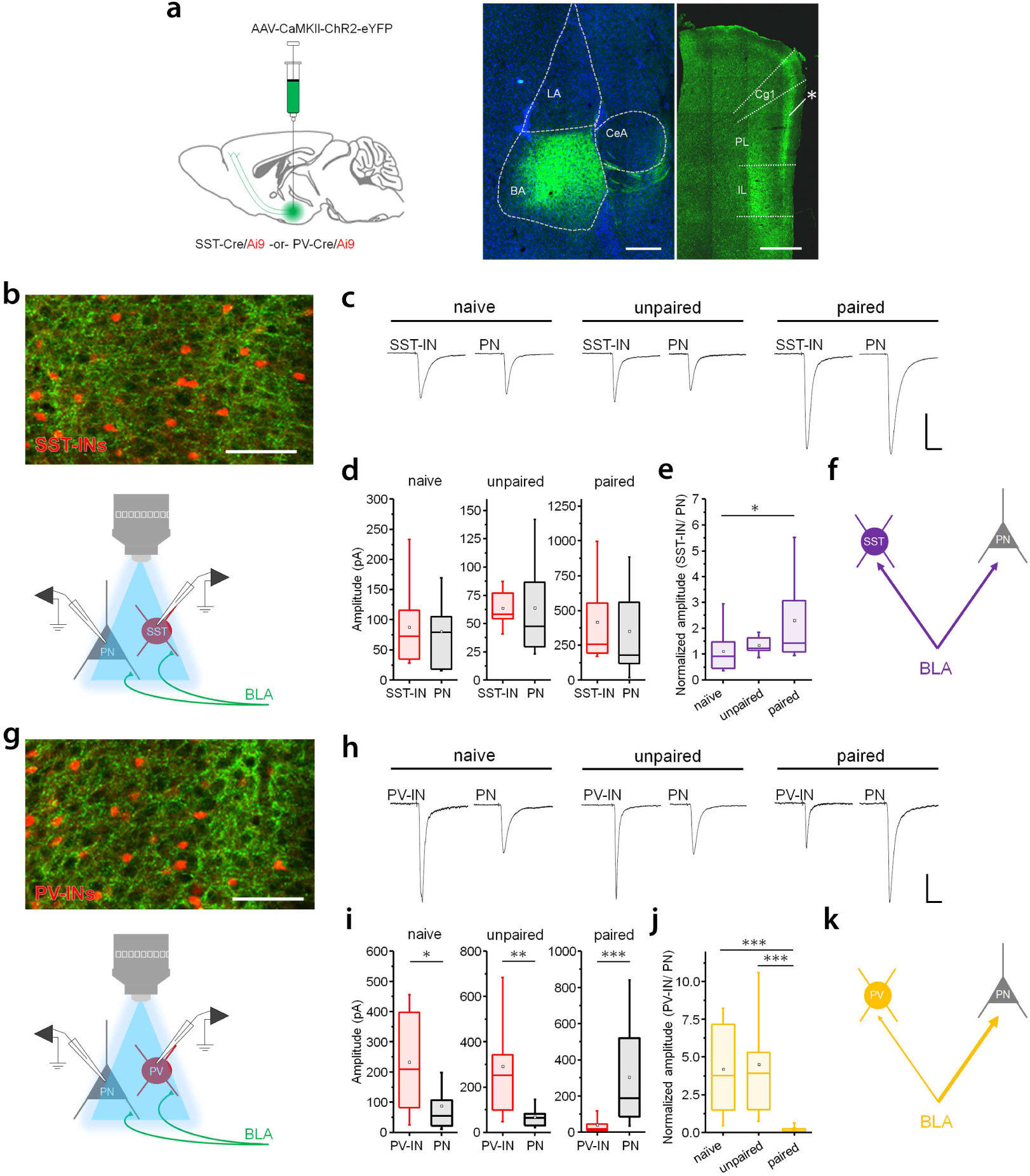
Balance of BLA transmission onto prelimbic INs and PNs is modulated by learning. **a,** To examine prelimbic (PL) neuronal responses to BLA afferent stimulation, a CaMKII promoter-dependent ChR2 vector was infused into the BLA (left) of SST- or PV-IRES-Cre/ Ai9 mice, leading to projection-specific expression of ChR2-eYFP in PL L2/3 (right). LA = lateral amygdala, BA = basal amygdala, CeA = central amygdala, Cg1 = cingulate area 1, IL = infralimbic. * PL ChR2 accumulation. Scale = 250 (left) and 500 (right) μm. At 24 hours after CS-US pairing, recordings were obtained from Tomato^+^ INs as well as surrounding PNs during BLA afferent photoexcitation (460 nm, 1 ms pulse, 0.1 Hz). **b**, Axonal ChR2 accumulation in layer 2/3 of SST-IRES-Cre/ Ai9 mice and recording configuration. **c**, Example EPSC traces. Scale = 100 pA x 50 ms. **d**, Amplitude of BLA-evoked SST-IN and PN EPSCs in slices from naïve mice (n = 4), as well as those that received unpaired (n = 3 mice) or paired training (n = 4 mice). Effect of cell type (naïve): *U* = 134, p = 0.51, Mann-Whitney *U* test; SST-IN, n = 18 cells; PN, n = 13 cells. Effect of cell type (unpaired): *U* = 106, p = 0.49, Mann-Whitney *U* test; SST-IN, n = 13 cells, PN, n = 14 cells. Effect of cell type (paired): *U* = 107, p = 0.37, Mann-Whitney *U* test; PV-IN, n = 16 cells; PN, n = 11 cells. **e**, Amplitude of EPSCs in SST-INs normalized to median values from PNs in the same slices. Effect of training: *χ*^2^ = 7.2 (2), p = 0.027, Kruskal-Wallis ANOVA. **f**, Relative strength of BLA transmission in conditioned mice. **g**, Axonal ChR2 accumulation in layer 2/3 of PV-IRES-Cre/ Ai9 mice. **h**, Example EPSC traces. Scale = 100 pA x 50 ms. **i**, Amplitude of BLA-evoked PV-IN and PN EPSCs in slices from naïve mice (n = 3), as well as those that received unpaired (n = 3 mice) or paired training (n = 4 mice). Effect of cell type (naïve): *U* = 118, p = 0.025, Mann-Whitney *U* test; PV-IN, n = 14 cells; PN, n = 11 cells. Effect of cell type (unpaired): *U* = 85, p = 0.0066, Mann-Whitney *U* test; PV-IN, n = 10 cells, PN, n = 10 cells. Effect of cell type (paired): *U* = 19, p = 7.56 × 10^−4^, Mann-Whitney *U* test; PV-IN, n = 13 cells; PN, n = 12 cells. **j**, Amplitude of EPSCs in PV-INs normalized to median values from PNs in the same slices. Effect of training: *χ*^2^ = 21.6 (2), p = 2.04 × 10^−5^, Kruskal-Wallis ANOVA. **k**, Relative strength of SST-IN transmission in conditioned mice. * p < 0.05, ** p < 0.01, *** p < 0.001 by Mann-Whitney *U* test (**i**) or Dunn’s post-hoc test (**e, j**). Box plots depict median (center line), mean (black box), quartiles, and 10-90% range (whiskers).

A requirement of cue-specific SST-IN activity during fear conditioning suggests that prefrontal SST-INs could mediate a critical component of memory encoding. However, previous studies have established that fear conditioning requires basolateral amygdala (BLA) activity ^15^, and memory storage is widely believed to be mediated by plasticity of synaptic connections within the lateral and basal nuclei ^12,13,16–22^. Therefore, because activation of prelimbic SST-INs is sufficient to evoke a fear response (Fig. 4c), we wondered whether deficits in learning after amygdala silencing can be explained in part by reduced prefrontal SST-IN transmission. To test this hypothesis, we performed photoinhibition of BLA excitatory neurons during CS-US pairing. One week prior to conditioning, SST-Cre x Ai9 mice received BLA injections of Arch or eYFP control vectors under the control of a CaMKII promoter (AAV-CaMKII-ArchT-GFP or AAV-CaMKII-eYFP) and were implanted with optic ferrules directed at the BLA (Fig. 5f; Supplementary Fig. 9). Fear conditioning and retrieval tests were conducted as described for prelimbic SST-IN photoinhibition (Fig. 5f). While freezing occurred in both groups during conditioning (Fig. 5g), Arch subjects exhibited a deficit in CS-evoked responses relative to eYFP subjects during the memory retrieval test (Fig. 5h). Following retrieval, whole-cell recordings revealed that SST-INs exhibited lower spontaneous EPSC frequency as well as increased paired-pulse ratios of evoked EPSCs (Fig. 5i-j) in Arch-expressing mice, compared to eYFP controls. These data independently confirm the relationship between memory encoding and increased SST-IN transmission, and suggest that this SST-IN plasticity depends at least in part on BLA activity.

**Fig. 9:**
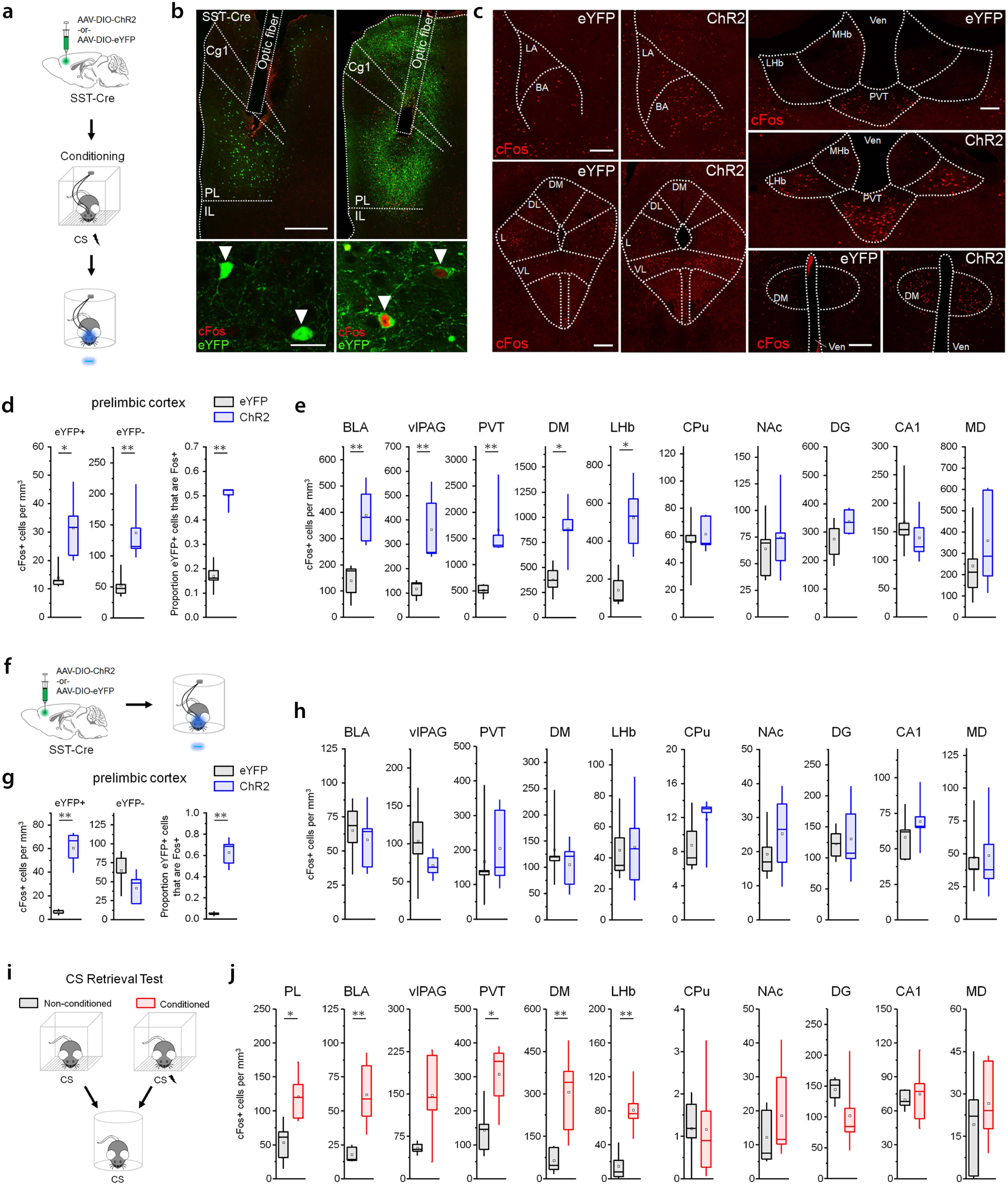
SST-IN activation recruits a specific brain network in conditioned mice. **a,** Preparation of ChR2 (n = 5) and eYFP mice (n = 5) used for cFos activity mapping after SST-IN photoexcitation. All subjects underwent CS-US pairing 24 hrs prior to stimulation. **b,** Induction of cFos by photoexcitation (6 trials, 473 nm, 10 ms pulses, 20 Hz) in prelimbic (PL) cortex. Insets (bottom) depict eYFP/ cFos double-labeled SST-INs. Arrowheads denote SST-INs. Scale = 500, 50 μm. **c**, Elevated cFos expression in select brain regions after optic stimulation of conditioned mice. LA = lateral, BA = basal, amygdala; DM = dorsomedial hypothalamus; DL = dorsolateral, L = lateral, VL = ventrolateral, periaqueductal gray; MHb = medial habenula; LHb = lateral habenula; PVT = paraventricular thalamus. Scale = 200 μm. **d,** Comparison of cFos^+^ cell counts in PL of stimulated ChR2-eYFP (blue) versus eYFP-only (black) mice. In both groups, eYFP+ cells represent transfected SST-INs, while eYFP-cells represent non-expressing neighbors. Group differences established by Mann-Whitney *U* test and controlled for false discovery (type 1 error) by the method of Benjamini and Hochberg. eYFP+ cells: *U* = 1, p = 0.016. eYFP-cells: *U* = 0, p = 0.0079. Proportion eYFP+ cells that are cFos+: *U* = 0, p = 0.0079. **e**, Comparison of cFos+ cell counts for all brain regions using the same statistical procedures as in (**c**). BLA: *U* = 0, p = 0.0079. vlPAG: *U* = 0, p 0.0079. PVT: *U* = 0, p = 0.0079. DM: *U* = 1, p = 0.0022. LHb: *U* = 0, p = 0.0012. Caudate putamen (CPu): *U* = 14, p = 0.84. Nucleus accumbens (NAc): *U* = 10, p = 0.69. Dentate gyrus of the dorsal hippocampus (DG): *U* = 7, p = 0.29. Cornu ammonis area 1 of the ventral hippocampus (CA1): *U* = 16, p = 0.55. Mediodorsal thalamus (MD): *U* = 8, p = 0.42. **f**, Preparation of ChR2 (n = 5) and eYFP mice (n = 5) used for cFos activity mapping after SST-IN photoexcitation in naïve (unconditioned) mice. **g**, Comparison of cFos^+^ cell counts in PL of stimulated ChR2-eYFP (blue) versus eYFP-only (black) mice, as conducted in (**c**). eYFP+ cells: *U* = 0, p = 0.0079. eYFP-cells: *U* = 20, p = 0.15. Proportion eYFP+ cells that are cFos+: *U* = 0, p = 0.0079. **h**, Comparison of cFos+ cell counts in select brain regions of stimulated naïve mice, as conducted for conditioned mice (**e**). BLA: *U* = 14, p = 0.84. vlPAG: *U* = 19, p 0.22. PVT: *U* = 11, p = 0.84. DM: *U* = 13, p = 1. LHb: *U* = 13, p = 1. CPu: *U* = 7, p = 0.31. NAc: *U* = 9, p = 0.55. DG: *U* = 12, p = 1. CA1: *U* = 6, p = 0.22. MD: *U* = 14, p = 0.84. **i**, Preparation of SST-IRES-Cre mice used for cFos activity mapping after CS exposure (4 trials, 2 KHz, 80 dB, 20 s) in mice that were conditioned 24 hours prior (n = 6) or in those that did not receive conditioning (n = 5). **j**, Comparison of cFos+ cell counts in select brain regions of conditioned versus nonconditioned mice, as conducted for photostimulated mice (**e** and **h**). PL: *U* = 2, p = 0.0017. BLA: *U* = 0, p = 0.0043. vlPAG: *U* = 5, p = 0.082. PVT: *U* = 3, p = 0.030. DM: *U* = 0, p = 0.0043. LHb: *U* = 0, p = 0.0043. CPu: *U* = 17, p = 0.79. NAc: *U* = 9, p = 0.33. DG: *U* = 25, p = 0.082. CA1: *U* = 12, p = 0.66. MD: *U* = 12, p = 0.66. * p < 0.05, ** p < 0.01 by Mann-Whitney *U* test. Box plots depict median (center line), mean (black box), quartiles, and 10-90% range (whiskers).

### Microcircuit organization and opposing behavioral roles of SST- and PV-INs

An increase in freezing upon SST-IN photoexcitation (Fig. 4c) is surprising given that SST-INs are likely to form inhibitory contacts onto excitatory PNs that control memory expression ^15^. However, we considered the possibility that another class of interneurons also receives input from SST-INs and, when inhibited via these connections, might be responsible for disinhibition of PNs ^23^. In particular, previous work indicates that mPFC PV-INs exhibit firing rate inhibition during CS exposure, and that photoinhibition of PV-INs elicits freezing ^4^. We therefore utilized optogenetics to establish whether SST-INs form functional connections with PV-INs, as well as to compare the strength of such connections with those formed onto PNs from the same brain slices. In addition, we asked whether CS-US pairing affects the balance of transmission from SST- or PV-INs onto inhibitory versus excitatory populations, which might modulate whether SST- or PV-IN recruitment leads predominantly to inhibition or disinhibition of PNs.

First, to enable selective interrogation of synaptic outputs from SST-INs, we microinjected an INTRSECT ChR2 vector (AAV-Cre_off_/Flp_on_-hChR2-eYFP) ^24^ into the prelimbic cortex of SST-IRES-FlpO/PV-IRES-Cre/Ai9 triple transgenic mice, permitting independent tagging of SST- (ChR2) and PV-INs (tdTomato) (Fig. 6a). We then examined inhibitory responses to SST-IN photoexcitation in PV-INs as well as PNs from the same brain slices to control for group differences attributable to viral expression. In both naïve and unpaired control mice, SST-INs elicited monosynaptic IPSCs in PV-INs that were less potent than those recorded in surrounding PNs (Fig. 6b-c; Supplementary Fig. 10). In contrast, after CS-US pairing, responses to SST-IN photoexcitation in these cell types were similar in amplitude. Comparison of SST-IN-evoked responses in PV-INs normalized to those in PNs confirmed that CS-US pairing increases the relative strength of SST-IN→PV-IN transmission (Fig. 6d). This suggests that learning shifts the balance of SST-IN output to favor inhibition of PV-INs, which may increase the likelihood of SST-IN-evoked disinhibition.

Next, we utilized a similar genetic approach to interrogate PV-IN transmission onto SST-INs and PNs in prelimbic layer 2/3 (Fig. 6f). Strikingly, this revealed that regardless of the training condition, PV-INs elicit IPSCs that are ~10-fold larger in amplitude in PNs compared to surrounding SST-INs (Fig. 6g-h). Comparison of SST-IN responses normalized to those in PNs revealed no effect of training on the balance of transmission (Fig. 6i). These results indicate the presence of a strong bias in PV-IN output that potentially favors preferential control of PN over SST-IN firing (Fig. 6j). In contrast, SST-INs exhibit a much weaker bias (~2-fold) for PNs over PV-INs, and this bias is completely eliminated by conditioning (Fig. 6e). This implies that SST-INs in the prelimbic cortex could have a unique capacity to evoke PN disinhibition.

Prelimbic circuit organization suggests that SST-INs might interact directly with PV-INs to mediate fear expression through PN disinhibition. As an *in vivo* test of this model, we first sought to confirm that photoinhibition of PV-INs elicits freezing, as reported previously ^4^. Indeed, after fear conditioning, PV-IN photoinhibition resulted in an increase in freezing during light-only trials (Fig. 7a, Supplementary Fig. 11). We then tested whether inhibition of PV-INs is required specifically for SST-IN-evoked freezing. In contrast to SST-IN-specific manipulations (Fig. 4c), concurrent photoactivation of SST- and PV-INs in SST-Cre/PV-Cre double transgenic mice was sufficient to negate the fear-promoting effect of SST-IN activity (Fig. 7b). These results imply that interaction between SST- and PV-INs is important for processing the behavioral output of SST-IN activity.

### BLA afferent connectivity of prelimbic INs

Having established that SST-INs directly inhibit PV-INs, we next considered whether these IN populations are engaged by long-range inputs to the prelimbic cortex. In addition to modulating plasticity of prelimbic SST-INs (Fig. 5), BLA contains prelimbic cortex-projecting neurons that exhibit increased firing during memory retrieval ^25,26^ and regulate fear memory expression ^27,28^. Projections from these cells primarily target layer 2/3 and are thus well-positioned to recruit potentiated SST-INs ^29^. To test whether prelimbic interneurons are directly modulated by these projections, we infused a calcium-calmodulin kinase II (CaMKII)-driven ChR2 vector (AAV-CaMKII-hChR2-eYFP; CaMKII-hChR2) into BLA, leading to axonal ChR2 accumulation in prelimbic layer 2/3 of PV- and SST-IRES-Cre/ Ai9 mice (Fig. 8a). Optic stimulation of these projections in naïve mice elicited compound EPSCs and feedforward IPSCs in both SST- and PV-INs, as well as surrounding PNs (Supplementary Fig. 12). Because much of this transmission occurred at long latencies after stimulation, recurrent activity within prelimbic circuits is likely responsible for its generation. Interestingly, compared to PNs from the same brain slices, SST-but not PV-INs exhibited a higher ratio of excitatory to inhibitory charge during these events (Supplementary Fig. 12c). This could be attributed to less potent network inhibition of SST-INs, since inhibitory charge in SST-INs was lower than in surrounding PNs (Supplementary Fig. 12b).

While the above results are intriguing, the complexity of BLA-evoked activity prohibited analysis of monosynaptic transmission at connections between BLA axons and prelimbic INs. We therefore applied a pharmacological cocktail to eliminate action potential propagation and prevent polysynaptic transmission ^12,13,30^. When using this approach in SST-Cre/ Ai9 mice, we found that regardless of training condition, BLA afferents evoke responses of similar amplitude in SST-INs compared to PNs (Fig. 8c-d; Supplementary Fig. 13). However, when SST-IN responses were normalized to those of PNs, this revealed a slightly higher ratio of SST-IN/ PN transmission in paired mice compared to naïve controls (Fig. 8e). In contrast to the above results, BLA terminal stimulation in naive and unpaired PV-Cre/ Ai9 mice evoked EPSCs that were larger in amplitude in PV-INs compared to PNs (Fig. 8h-j). This is in agreement with similar experiments that examined the potency of BLA transmission onto PV-INs in the infralimbic cortex ^31^. However, in animals that received CS-US pairing the relative strength of BLA transmission in PV-INs and PNs was effectively reversed (Fig. 8j). These data collectively imply that, following conditioning, SST-INs are likely strongly activated by BLA afferents and that circuit plasticity may favor their recruitment over PV-INs (Fig. 8f, k).

### Network disinhibition underlies SST-IN-evoked fear expression

Because the balance of ongoing excitatory and inhibitory transmission determines the firing rate of excitatory PNs, the activity of prelimbic output neurons could be modulated solely through the relief of PV-IN-mediated inhibition. To reveal the extent to which SST-IN activity disinhibits prelimbic networks, we therefore conducted immunolabeling for the activity marker cFos in prelimbic cortex and potential downstream brain regions following SST-IN photoexcitation, in the absence of any CS exposure, at 24 hours after conditioning (Fig. 9a; Supplementary Fig. 14). Similar to a previous experiment (Fig. 4), ChR2-expressing mice but not eYFP controls exhibited an increase in freezing in response to photoexcitation (Supplementary Fig. 14). After behavioral testing, ChR2 mice exhibited higher cFos labeling of SST-INs compared to eYFP controls, as well as higher cFos labeling of surrounding eYFP-negative cells, consistent with disinhibition of other prelimbic cell types (Fig. 9b, d). When quantification was extended to downstream targets of prelimbic cortex, higher numbers of cFos-positive cells were also detected in the BLA, paraventricular thalamus, lateral habenula, ventrolateral periaqueductal gray, and the dorsomedial hypothalamus (Fig. 9c, e). However, several other regions including the nucleus accumbens, caudate putamen, ventral hippocampus area CA1, dentate gyrus, and mediodorsal thalamus were unaffected by photostimulation (Fig. 9e; Supplementary Fig. 14).

To test whether regional cFos induction by SST-IN photoexcitation occurs independently of fear expression, we next quantified cFos expression in a randomly selected subset of naïve mice that received optogenetic manipulation of prelimbic SST-INs without prior fear conditioning (Fig. 4f). Consistent with the larger group, these mice exhibited no increase in freezing over baseline levels during photostimulation (Fig. 9f; Supplementary Fig. 15). Examination of stimulated prelimbic tissue confirmed that consistent with photoactivation, higher cFos labeling was present in eYFP-positive cells of ChR2-relative to eYFP-expressing mice (Fig. 9g; Supplementary Fig. 15). However, there was no group difference in cFos expression in surrounding eYFP-negative cells, indicating that in contrast to animals that received CS-US pairing (Fig. 9b, d), SST-IN photoexcitation in naïve mice does not activate surrounding prelimbic neurons to a significant degree. In addition, remote brain regions that were modulated by photoexcitation in conditioned mice did not exhibit any group differences in the number of cFos-positive neurons (Fig. 9h; Supplementary Fig. 15). Thus, acquisition of SST-IN-evoked freezing correlates with a change in SST-IN recruitment of a specific brain network including prelimbic neurons indirectly activated by SST-IN photoexcitation, presumably via disinhibition.

Finally, to test whether network-level effects of memory retrieval resemble those evoked by SST-IN photoexcitation in conditioned mice, we performed cFos analysis following CS exposure (Fig. 9i). Presentation of 4 CS trials elicited increased freezing in mice that received CS-US pairing 24 hours prior to the memory retrieval test, but not in non-conditioned controls (Supplementary Fig. 16). After CS exposure, a higher number of cFos-positive cells was observed in conditioned relative to non-conditioned mice in the majority (5/6) of brain regions that were modulated by SST-IN photoexcitation (Fig. 9j). The remaining region (vlPAG) showed a trend toward higher cFos labeling in conditioned mice (p = 0.082). Conversely, areas in which cFos immunoreactivity was unaffected by SST-IN photoexcitation also exhibited no differences in cFos-positive cells following CS-evoked memory retrieval. Together these data argue against the notion that network cFos induction by photostimulation results from nonphysiological activity patterns, and suggest that CS recruitment of SST-INs mediates disinhibition of prelimbic outputs to remote brain regions underlying memory expression.

## Discussion

In this study we demonstrate that associative fear conditioning potentiates the function of prefrontal SST-INs at the level of synaptic transmission and *in vivo* activity. Correlated with memory acquisition, SST-INs exhibit an experience-dependent increase in CS-evoked firing and specifically signal the learned CS rather than the general fear state of the animal. Activation of SST-INs in turn plays a causal role in both the expression as well as initial acquisition of conditioned fear responses. The paradoxical role of SST-INs in memory expression can be explained by their encoding of cue-related disinhibition of prelimbic PNs, which are in turn responsible for the recruitment of a distributed brain network for defensive responding.

In network disinhibition, prelimbic circuits formed by SST- and PV-INs exhibit functional differences that may facilitate their complementary roles. In particular, output from PV-INs is heavily biased toward PNs over SST-INs (Fig. 6), implying that they are specialized for suppression of PN firing. In contrast, there is a greater potential for PN disinhibition during SST-IN activity, owing to their relatively strong inhibition of PV-INs. Accordingly, photoactivation of SST-INs indirectly activates not only surrounding prelimbic cells but also remote brain regions that receive excitatory connections from the mPFC (Fig. 9). A learning-dependent shift in SST-IN output could in part explain why these effects are specific to conditioned mice. Alternatively, plasticity in other circuits could prime this network for SST-IN-evoked disinhibition.

Although all prelimbic cell types that we examined receive direct input from BLA projections, it is notable that these afferents evoke complex synaptic activity with an overall higher ratio of excitatory: inhibitory transmission in SST-INs (Supplementary Fig. 12). In conditioned mice, PV-INs but not SST-INs exhibited a lower strength of monosynaptic BLA transmission compared to neighboring PNs (Fig. 8). Given these findings, it seems possible that after learning BLA afferent activity favors the recruitment of SST-INs over PV-INs, and that resulting PN disinhibition could underlie freezing behavior elicited by BLA afferent stimulation in conditioned mice ^27^. In addition, BLA projections could also mediate the effect of amygdala activity on learning-related SST-IN plasticity (Fig. 5). Further interrogation of synaptic and functional plasticity throughout this circuitry will be necessary to establish the degree to which other cell types, and projection-specific populations, cooperate with SST-INs in memory encoding and retrieval.

An important implication of SST-IN plasticity is that phasic disinhibition, which has been established primarily as a mechanism for cortical processing of behavioral feedback signals ^23,32–35^, can be encoded by potentiation of GABAergic transmission to mediate novel cue associations. This means that both activated (e.g. SST-INs) and inhibited interneuron populations (e.g. PV-INs) can play a role in memory expression and exhibit cue-evoked activity patterns that are causally linked. In this way the modification of a sparse interneuron population can extensively reorganize stimulus processing in a broader local network. Indeed, previous findings confirm that firing rate inhibition of PV-INs is a potent regulator of PN recruitment, synchrony, and synaptic plasticity ^4,8,23,32,35^. Nevertheless, it remains unclear why SST-IN-evoked disinhibition triggers such selective changes in behavior and circuit activity. One possibility is that different subnetworks of prelimbic PNs, for example populations with discrete projection patterns, vary in the level of inhibition they receive from SST-versus PV-INs. The balance of ongoing activity from these inhibitory sources could in part determine whether a given cell is predominantly activated, inhibited or unaffected by cue-responsive SST-INs.

Several brain regions activated by prelimbic SST-IN stimulation, including the BLA ^15^, paraventricular thalamus ^36,37^, lateral habenula ^38^, ventrolateral periaqueductal gray ^39^, and the dorsomedial hypothalamus ^40^, have established roles in mediating defensive behavioral responses to conditioned and unconditioned stimuli. It is possible that PNs that give rise to direct projections to these regions from the prelimbic cortex are preferentially disinhibited during SST-IN firing. Alternatively, this pattern of network activity may be completed by relay circuits from other downstream effectors. Intriguingly, however, neither the striatum nor the mediodorsal thalamus, which are among the areas receiving the highest density of input from the prelimbic cortex ^41,42^, are activated during the expression of SST-IN-evoked freezing. This implies a high level of specificity in the computational logic of prelimbic microcircuits.

While our *in vivo* manipulations were targeted to SST-INs as a whole, somatostatin expression demarcates a cell population with heterogeneous protein expression, firing patterns and morphology ^43^, whose function is additionally determined by their laminar location ^44^. Our results imply that SST-INs that encode fear-related disinhibition reside in superficial layers of the prelimbic cortex, but further parcellation of this cell class could reveal functionally discrete subpopulations. For example, previous work has identified a subset of SST-INs that express the oxytocin receptor and contribute to the regulation of anxiety and social behavior in a sex-dependent manner ^45,46^. At present it remains unclear what features of an aversive experience might be encoded by SST-INs or how this information is represented by specific SST-IN populations. Future studies could resolve the function of individual neurons through the use of genetic markers, cellular tags and *in vivo* imaging. In addition, although circuit disinhibition can be explained by fast GABAergic transmission, it is also important to consider that somatostatin is not just a marker of interneurons but also a peptide transmitter that can influence memory acquisition ^47^ and recall ^48^. The release of this peptide from dense-core vesicles may contribute to the memory function of potentiated SST-INs.

In conclusion, our results outline an important casual role for inhibitory signaling in associative memory. In pursuit of memory engrams, it will therefore be critical to consider the contributions of interneurons that, despite their inhibitory output, have the capacity to encode and reactivate a specific pattern of excitatory neuronal activity. Interrogation of these cells and their associated circuitry could reveal important computational principles for the storage, consolidation and retrieval of information. Indeed, during the preparation of our revised manuscript, a new report demonstrated an important causal role for prefrontal SST-IN activity in learned social fear ^49^.

## Acknowledgments

We thank Sabina Bayshtok and Emily Beckett for expert technical assistance, behavioral scoring and assistance with experimenter blinding; Ciorana Roman-Ortiz for help with optogenetic behavioral manipulations; and Denise Cai for comments on the manuscript. These experiments were supported by funds from National Institute of Mental Health (NIMH) grants RO1 MH105414, RO1 MH116145, and R21 MH114170 to R.L.C, in addition to F32 MH115688 to K.A.C.

## Author contributions

K.A.C. and R.L.C. initiated the project; R.L.C. supervised research; K.A.C. and R.L.C. designed experiments; K.A.C. performed the research and data analysis; R.L.C. and K.A.C. wrote the manuscript.

## List of Supplementary Materials

Methods

Supplementary Figs. 1-16

