## Supplementary Materials for "Prefrontal somatostatin interneurons encode fear memory"

**Supplementary Methods**

*Animals*

All experimental procedures were approved by the Institutional Animal Care and Use Committee at the Icahn School of Medicine at Mount Sinai. Experiments were performed on male mice aged P42-P60. Mice were acquired from Jackson Laboratories (Bar Harbor, ME, USA) and maintained in the C57Bl/6J background. The following genotypes were used: SST-IRES-Cre (Stock No. 028864), PV-IRES-Cre (Stock No. 017320), SST-IRES-FlpO (Stock No. 028579), PV-IRES-FlpO (Stock No. 022730), and Ai9 (Stock No. 007909). Mice were housed 2-5 per cage in a 12 hr light-dark cycle with access to food and water *ad libitum*.

*Stereotaxic vector infusion and optic fiber implantation for in vivo optogenetics*

Viral constructs were purchased from University of Pennsylvania Vector Core or Addgene and included AAV1-EF1a-DIO-hChR2(H134R)-eYFP-WPRE (Addgene # 20298), AAV1-CBA-FLEX-Arch-GFP (Addgene # 22222), AAV1-EF1a-DIO-eYFP-WPRE (Addgene # 27056), AAV1-CaMKIIα-ArchT-GFP (Addgene # 99039), and AAV1-CaMKIIα-eYFP-WPRE (Addgene # 105622)Optic fibers (200 μm diameter, Thor labs) were fixed inside ferrules (1.25 mm OD, 230 μm ID, Precision Fiber Products, Chula Vista, CA, USA) using heat-cured epoxy, cut to 2.5 mm length for mPFC or 5.0 mm length for basolateral amygdala, and polished using aluminum oxide lapping paper with incremental decreases in graininess (Thor labs, Newton, NJ, USA). Light transmittance was measured using a light intensity meter (Thor labs) at wavelengths used in each respective experiment. Stereotaxic surgeries were conducted at P45-49. Anesthesia was induced using 5% inhaled isoflurane vaporized in oxygen delivered at a rate of 1-1.5 L/min. Mice were then mounted in stereotaxic frames and maintained at 1-1.5% isoflurane (Stoelting, Wood Dale, IL, USA or Kopf, Tujunga, CA). Viral constructs were injected bilaterally into prelimbic cortex (300 nL; AP +1.9, DV -2.0, ML +/- 0.9 at a 10° angle) or basolateral amygdala (250 nl; AP

-1.4, DV -5.1, ML +/- 3.3) using motorized injectors (Stoelting and World Precision Instruments) at a rate of 100 nL/min. Following infusion, the needle was left in place for an additional 10 minutes prior to slow removal at a rate of 0.03 mm/second. For *in vivo* optogenetics, following bilateral injections, optic ferrules were bilaterally implanted and directed towards prelimbic cortex (AP +1.9, DV -1.6, ML +/- 0.9 at a 10° angle) or basolateral amygdala ( AP -1.4, DV -4.8, ML +/- 3.3). Ferrules were fixed in place using C&B Metabond luting cement (Parkell, Edgewood, NY, USA) and dental cement. Post-surgical analgesia was achieved with banamine (2.5 mg/kg). Mice recovered in home cages for at least 1 week prior to experimental manipulation.

*Stereotaxic vector infusion for electrophysiology*

AAV1-CaMKIIα-hChR2(H134R)-eYFP-WPRE (Addgene # 26969P) was purchased from the University of Pennsylvania Vector Core. rAAVDJ/nEF-$Cre_{off}$/$Flp_{on}$-hChR2(H134R)-eYFP was purchased from the University of North Carolina Gene Therapy Center Vector Core. Stereotaxic surgeries were done at P42-45 for AAV1-CaMKIIα-hChR2(H134R)-eYFP-WPRE and from P25-P28 for AAVDJ-$Cre_{off}$/$Flp_{on}$-hChR2(H134R)-eYFP. AAV1-CaMKIIα-hChR2(H134R)-eYFP-WPRE and rAAVDJ/nEF-$Cre_{off}$/$Flp_{on}$-hChR2(H134R)-eYFP were bilaterally infused into basolateral amygdala (250 nl; AP -1.3, DV -5.1, ML +/- 3.3) or prelimbic cortex (400 nL; AP +1.9, DV -1.6, ML +/- 0.2), respectively. Mice recovered for 10 days (AAV1-CaMKIIα-eYFP-WPRE) or 25-27 days ($rAAV_{DJ}$/nEF-$Cre_{off}$/$Flp_{on}$-hChR2(H134R)-eYFP) prior to electrophysiological recordings, which we determined to be the incubation time required for effective expression of ChR2 by these viral serotypes.

*Optogenetic behavioral manipulations*

Mice were acclimated to handling and patch cord tethering for 3 consecutive days prior to behavioral testing. Handling consisted of 10 min handling sessions, followed by 15 min habituation to patch cords in a fresh cage, followed by 5 min of additional handling. Fear conditioning and retrieval tests were conducted in sound attenuating chambers with automated stimulus delivery software (MedAssociates, St. Albans, VT, USA). Each session began with a 200 s baseline period after which various stimuli were presented with a fixed 80 s interstimulus interval. Auditory fear conditioning entailed 6 pairings of an auditory tone (CS; 2kHz, 80 dB, 20 s) with a co-terminating footshock (US; 0.7 mA, 2 s). For experiments where optogenetic manipulations were performed during memory retrieval, animals were conditioned while connected to patch cords but no light stimuli were delivered. Twenty-four hours after conditioning, modulation of freezing behavior by CS and light stimuli were examined in a context distinct from the conditioning arena (context B). In the retrieval test, mice were subjected to two laser-only trials, followed by 4 CS presentations alternating with and without concurrent laser stimulation. CS-only or CS + light presentation order was counterbalanced. The two laser-only trials, two CS-only, and two CS+light trials were each averaged and used for subsequent analysis. In experiments in naïve mice, animals were subjected to two trials of laser stimulation in context B without prior fear conditioning. For trials including optogenetic stimulation, a final transmitted intensity of 7-9 mW for 473 nm (for ChR2; 20 Hz, 10 ms for 20 s epochs) or 563 nm (for Arch; constant light, 20 s) laser-generated (Opto Engine, LLC, Midvale, UT, USA) light was used. Parameters for photoexcitation are consistent with other recent studies of cortical SST-INs (*1, 2*), and the 20 Hz firing rate observed during *ex vivo* stimulation of SST-INs (Supplementary Fig. 6) is within the endogenous range of activity in prelimbic SST-INs (*3*). In experiments where neurons were silenced during learning, light was presented simultaneously with each CS and extended 3 seconds after CS/US termination, after which it was manually ramped down to 0 mW over a period of 3 seconds. Twenty-four hours later, mice were tethered to patch cords in context B and subjected to 4 CS presentations in the absence of light stimulation. Mice were then used for electrophysiology recordings 24 hours after retrieval. Behavior was recorded by video and scored offline by a trained experimenter blind to the identity of the subject. Scoring by a second blinded experimenter was used to validate results. Fiber tip placement and viral expression were confirmed by fluorescence microscopy. Subjects having misplaced fiber tips or lacking viral expression encompassing at least ~50% of the target structure, in either hemisphere, were excluded from analysis by an investigator blind to experimental results.

*Open field*

Naïve SST-IRES-Cre mice expressing ChR2 or eYFP in prefrontal SST-INs were used for open field experiments. Mice were habituated to the room for 30 minutes prior to the start of the experiment. Mice were tethered to patch cords and placed into the center of a 42 cm (length) x 42 cm (width) x 30 cm (height) square arena equipped with 15 infrared beams/detectors on each side. Tests were 20 min long and consisted of alternating light on (473 nm, 7-9 mW delivered at 20 Hz, 10 ms pulses) and light off periods lasting 5 minutes each. Light on/off epochs were counterbalanced in both groups. Open field arenas were connected to a PC running Fusion v5.6 SuperFlex software, which was used to analyze infrared beam breaks and quantify locomotor parameters. Locomotor metrics are reported as an average of each of the two light on or two light off periods.

*Fiber photometry surgery and calcium imaging*

SST-IRES-Cre mice received unilateral infusion (400 nl; A/P: +1.9, D/V: -1.6, M/L: + or - 0.2) of AAV1-hSyn-FLEX-GCaMP6f-WPRE (Addgene # 100833). Following viral infusion, imaging fibers fabricated from 400 µm core 0.48 numerical aperture optic fiber fixed in a 2.5 mm diameter metal ferrule (Doric Lenses, 2.0 mm length) were chronically implanted above the injection site in prelimbic cortex (A/P: +1.9, D/V: -1.5, M/L: + or - 0.2). Fibers were fixed to the skull using luting (Metabond) and dental cement. Mice were returned to their home cage for 4 weeks prior to the start of imaging, which was determined to be the amount of time required to achieve maximal GCaMP6f expression in prefrontal SST-INs.

Mice were habituated to handling (10 min) and patch cord tethering (15 min) for 3 consecutive days prior to the start of imaging experiments. Additionally, calcium signals were streamed during handling and habituation sessions to ensure robust and reproducible signals prior to the start of experiments. Mice were connected to an imaging patch cord (400 µm core; 0.48 numerical aperture, Doric Lenses), which was connected to a 6-port fluorescence minicube (Doric lenses). Blue (465 nm for GCaMP6f excitation, Doric lenses) and violet (405 nm for control artifact fluorescence, Doric Lenses) light was transmitted into the brain at 20-80 µW and kept constant across experimental sessions. Emitted light was passed through a dichroic mirror and a 500-540 nm filter before detection by a visible femtowatt photoreceiver (model 2151, Newport). Analog signals were recorded using a RZ5 processor and a PC equipped with Synapse software (Tucker-Davis Technologies). All conditioning was performed in MedAssociates operant chambers. For paired fear conditioning experiments, mice were tethered to a patch cord and exposed to 4 CS tones (2kHz, 80 dB, 20 s) during two test sessions at 24 hours pre- and post-conditioning. Paired conditioning consisted of 6 pairings of CS with a co-terminating US (0.7 mA, 2 s).

For unpaired conditioning, mice were tethered to the patch cord for both CS and US presentations but were untethered and returned to the home cage for 15 minutes between sessions. After 24 and 48 hours, mice were tethered to the patch cord and exposed to context B and context A, respectively. Calcium signals were sampled at 6kHz and were continuously recorded throughout all tests, including conditioning, CS, and context memory retrieval experiments. All imaging sessions started with a 2 minute baseline period during which calcium signals were recorded prior to the start of the experiment. The start and end of each experimental session as well as CS and US onset and offset timestamps were generated using TTL signals triggered by MedAssociates software to enable precise temporal analysis of calcium signals.

*Fiber photometry data analysis*

Extraction and analysis of fiber photometry signals was performed using custom code in MATLAB R2018b (original code available on Tucker-Davis Technologies website), following conventional practices reported elsewhere (*4*-*6*). Demodulated 465 nm and 405 nm signals were digitally filtered and scaled. To account for artifacts resulting from movement, photobleaching, and autofluorescence, which are reflected by fluctuations in the control 405 nm trace, the control signal was subtracted from the GCaMP6f signal (465 nm). The resulting fitted trace was then normalized to the 405 nm trace ($\Delta F/F$ = (465 nm signal – fitted 405 nm signal)/fitted 405 nm signal) and used to calculate changes in fluorescence during behavior. Changes in peak fluorescence in response to a 20-s CS or 2-s US presentation were calculated by normalization to a baseline period of equal duration occurring immediately prior to CS or US onset, respectively. For analysis of inter-CS freezing and contextual freezing epochs, behavior videos were scored to extract freezing onset and offset. Calcium traces associated with the freezing epochs were then aligned and averaged. Changes in fluorescence during the first 2 seconds of freezing, which was the minimum duration of freezing bouts, were compared to a 2 second pre-freezing baseline period immediately preceding the freezing epoch. Percent change in signal (%$\Delta F/F$) was calculated by subtracting the peak fluorescence signal in the test (CS, US, or freezing epoch) period from the peak fluorescence signal in the respective baseline period and dividing the resulting signal by the peak fluorescence signal in the baseline period (%$\Delta F/F = (F_{peakTEST} - F_{peakBL})/F_{peakBL}$). For frequency analysis of calcium-related events, the median absolute deviation of the fitted signal trace was calculated and all transients that exceeded 2.91 deviations were manually counted.

*Fear conditioning for electrophysiology*

Cued auditory fear conditioning entailed 6 pairings of an auditory tone (CS; 2kHz, 80 dB, 20 s) with a co-terminating footshock (US; 1 mA, 2 s). Control and experimental subjects were selected from the same litter and recordings from these groups were interleaved. Naive controls received only cage experience, while unpaired controls were subjected to 6 CS presentations followed 15 min later by 6 US presentations. Mice were sacrificed for electrophysiological analyses at 24 hours after conditioning.

*Slice electrophysiology*

Mice were deeply anesthetized using isoflurane inhalation prior to decapitation. Acute brain slices were prepared from medial prefrontal cortex at 350 μm thickness on a VT1200S vibratome (Leica Microsystems, Buffalo Grove, IL, USA) in a low sodium sucrose solution bubbled with carbogen (95% $O_2$, 5% $CO_2$) and consisting of (in mM): 210 sucrose, 26.2 $NaHCO_3$, 11 glucose, 2.5 KCl, 1 $NaH_2PO_4$, 0.5 ascorbate, 4 $MgCl_2$, and 0.5 $CaCl_2$, and chilled to -3-4°C. Slices were transferred to a recovery chamber continuously bubbled with carbogen and containing normal artificial cerebrospinal fluid (ACSF) consisting of (in mM): 119 NaCl, 26.2 $NaHCO_3$, 11 glucose, 2.5 KCl, 1 $NaH_2PO_4$, 2 $MgCl_2$, and 2 $CaCl_2$, and warmed to 34°C for 45 min. Following recovery, slices were maintained at room temperature until initiating recordings. Whole-cell electrodes were pulled from borosilicate glass and filled with a low chloride solution (for voltage clamp recordings) consisting of (in mM): 120 Cs-methanesulfonate, 10 HEPES, 10 Na-phosphocreatine, 8 NaCl, 1 QX-314, 0.5 EGTA, 4 Mg-ATP, and 0.4 Na-GTP or a K-based solution (for current clamp recordings) consisting of (in mM): 127.5 K-methanesulfonate, 10 HEPES, 5 KCl, 5 Na-phosphocreatine, 2 $MgCl_2$, 0.6 EGTA, 2 Mg-ATP, and 0.3 Na-GTP. Internal solutions were adjusted to pH 7.25 and 290-300 mOsm. Slices were visualized on an upright differential interference contrast microscope and LED-coupled (Prizmatix, Givat-Shmuel,

Israel) 40X objectives were used for the identification of fluorescently-tagged cells as well as optogenetic stimulation. SST- and PV-INs were identified based on tdTomato or EYFP fluorescence, with high membrane resistance (>100 MΩ) or low capacitance (< 90 pF) as confirmation of interneuron identity. Principal excitatory projection neurons (PNs) were identified based on morphology (large pyramidal soma and prominent apical dendrite), with low membrane resistance (<75 MΩ) or high capacitance (>100 pF) as additional criteria. Spontaneous excitatory (EPSCs) and inhibitory (IPSCs) postsynaptic currents were isolated by clamping neurons at -60 or 0 mV, respectively, in low-chloride internal solution. A total trace duration of at least 5 minutes was sampled at each potential for spontaneous currents. For paired-pulse measurements, EPSCs were evoked with a bipolar stimulating electrode placed in layer 2 of prelimbic cortex adjacent to the targeted postsynaptic neuron. To stimulate light-evoked transmission, we used TTL-pulsed microscope objective-coupled light-emitting diodes (LEDs, 460 nm, 20 mW/mm$^2$, Prizmatix). This intensity evoked maximal response amplitudes at a pulse duration of 1 ms. Spontaneous postsynaptic currents (Fig. 1 and fig. S1), paired-pulse analysis (Fig. 1) and light-evoked compound currents (fig. S4) were conducted in standard ACSF. For more stringent isolation of monosynaptic currents during optic stimulation (Fig. 3), recordings were conducted in the presence of 1 µm tetrodotoxin (Abcam) and 100 µm 4-aminopyrimidine (Abcam), which results in complete elimination of polysynaptic activity (*27, 28*).

Data were low-pass filtered at 3kHz (evoked) or 10 kHz (spontaneous) and acquired at 10 kHz using Multiclamp 700B (Molecular Devices, San Jose, CA, USA) and pClamp 10 software (Molecular Devices). Experimenter was blind to cell type and experimental condition during analysis of evoked (Clampfit 10, Molecular Devices) and spontaneous currents (MiniAnalysis, Synaptosoft, Fort Lee, NJ, USA).

*cFos immunofluorescence*

For analysis of CS-evoked cFos (Fig. 3), mice were subjected to 4 CS presentations in context B. For analysis of cFos induction by SST-IN photoexcitation (Fig. 9), ChR2 or eYFP-expressing mice were subjected to 6 bouts of 20 s stimulation (473 nm, 10 ms pulse, 20 Hz). Ninety minutes following CS or light stimuli, mice were deeply anesthetized via isoflurane inhalation and transcardially perfused with phosphate buffered saline (PBS) followed by 4% paraformaldehyde (PFA) in PBS (pH 7.45). Brains were post-fixed in PFA for 14-16 hours post perfusion and sectioned in 50 $\mu$m thick slices on the coronal plane on a VT1000S vibratome (Leica). Immunofluorescence staining against cFos was conducted on floating sections using a rabbit anti-cFos primary antibody (1:10000, Sigma Aldrich, F7799). Fluorescence-conjugated secondary antibodies included goat anti-rabbit conjugated to FITC (1:500, 111-095-003) and goat anti-rabbit conjugated to Alexa 647 (1:500, 111-605-003) and were purchased from Jackson Immunoresearch (West Grove, PA, USA). Slices were blocked in 2% goat serum in 0.3% Tween-80 PBS for 1 hour at room temperature. The primary antibody was incubated overnight at 4°C in 2% goat serum in 0.3% Tween-20 PBS. Following PBS washes, slices were incubated with the secondary antibody in 2% goat serum in 0.3% Tween-80 for 2 hours at room temperature. Following additional PBS washes, slices were incubated for 7 minutes at room temperature with filtered 1 mg/mL DAPI solution, followed by thorough PBS washes. Slices were then mounted with Prolong Antifade Gold mountant medium (Life Technologies, Grand Island, NY, USA) and imaged on a Zeiss confocal microscope operating Zeiss Zen software (Carl Zeiss Microscopy, Jena, Germany). tdTomato+ and eYFP+ SST-INs, as well as cFos+ nuclei, were manually quantified using the Cell Counter plugin in ImageJ (National Institutes of Health, Bethesda, MD, USA) while blind to experimental condition.

*Statistical analysis*

Prior to parametric statistical analysis, the Shapiro-Wilk test and Levene's test were used to establish normality of data and homogeneity of variance, respectively. Failing these assumptions or in cases in which group sizes might be too small to establish normality, we utilized non-parametric statistical comparisons. All figures utilizing bar graphs contain individual sample data, means, and standard error bars. Parameters used for box plots are included in corresponding figure legends. For reported effects, statistical power exceeded a minimum of 0.8, and in most cases 0.9. P-values obtained from network analysis of cFos+ cells were corrected for a false discovery rate of 10% using the method of Benjamini and Hochberg. Statistical analysis and graphing were conducted in Graphpad Prism (San Diego, CA) and OriginPRO (OriginLab, Northampton, MA).

### Supplementary Figure Legends

**Supplementary Figure 1. Spontaneous synaptic transmission in prelimbic interneurons after fear conditioning.** Example raw traces, mean interevent interval (IEI) and amplitude of events are shown. Whole-cell recordings were obtained from acute brain slices from naïve mice or littermates that were subjected paired or unpaired auditory fear conditioning, as in Fig. 1. Scale for all traces = 20 pA x 1 s. **a,** CS-evoked freezing during fear conditioning for SST-IRES-Cre/ Ai9 mice used for spontaneous transmission analysis. **b**, L2/3 SST-IN sEPSC amplitude, effect of training: $F_{2,31} = 0.58$, $p = 0.56$, 1-way ANOVA; naïve, n = 11 cells (3 mice); unpaired, n =12 cells (3 mice); paired, n = 11 cells (4 mice). **c**, L2/3 SST-IN sIPSCs. Interevent interval, effect of training: $\chi^2 = 1.35$ (2), $p = 0.51$, Kruskal-Wallis ANOVA; sIPSC amplitude, effect of training: $F_{2,31} = 1.18$, $p = 0.32$, 1-way ANOVA; naïve, n = 11 cells (3 mice); unpaired, n =12 cells (3 mice); paired, n = 11 cells (4 mice). **d**, L5/6 SST-IN sEPSCs. Interevent interval, effect of training: $F_{2,24} = 4.45$, $p = 0.023$, 1-way ANOVA; amplitude: $F_{2,24} = 0.12$, $p = 0.89$, 1-way ANOVA; naïve, n = 8 cells (3 mice); unpaired, n = 8 cells (3 mice); paired, n = 11 cells (3 mice). **e**, L5/6 SST-IN sIPSCs. Interevent interval, effect of training: $F_{2,19} = 0.12$, $p = 0.89$, 1-way ANOVA; amplitude, effect of training: $F_{2,19} = 0.15$, $p = 0.86$, 1-way ANOVA; naïve, n = 6 cells (3 mice); unpaired, n = 5 cells (3 mice); paired, n = 11 cells (3 mice). **f**, CS-evoked freezing during fear conditioning for PV-IRES-Cre/ Ai9 mice used for spontaneous transmission analysis. **g**, L2/3 PV-IN sEPSC amplitude, effect of training: $F_{2,36} = 0.45$, $p = 0.64$, 1-way ANOVA; naïve, n = 13 cells (3 mice); unpaired, n =13 cells (3 mice); paired, n = 13 cells (3 mice). **h**, L2/3 PV-IN sIPSCs. Interevent interval, effect of training: $F_{2,31} = 0.23$, $p = 0.80$, 1-way ANOVA; amplitude: $F_{2,31} = 0.22$, $p = 0.80$, 1-way ANOVA; naïve, n = 12 cells (3 mice); unpaired, n = 9 cells (3 mice); paired, n = 13 cells (3 mice). **i**, L5/6 PV-IN sEPSCs. Interevent interval, effect of training: $F_{2,18} = 1.04$, $p = 0.37$, 1-way ANOVA; amplitude, effect of training: $F_{2,18} = 5.41$, $p = 0.014$, 1-way ANOVA; naïve, n = 8 cells (3 animals); unpaired, n = 6 cells (3 animals); paired, n = 7 cells (3 animals). **j**, L5/6 PV-IN sIPSCs. Interevent interval, effect of training: $F_{2,18} = 1.83$, $p = 0.19$, 1-way ANOVA; amplitude, effect of training: $F_{2,18} = 2.48$, $p = 0.11$, 1-way ANOVA; naïve, n = 8 cells (3 animals); unpaired, n = 6 cells (3 animals); n = 7 cells (3 animals). * $p < 0.05$ by Tukey's post-hoc test (**d**) or Dunn's post-hoc test (**i**). Bar graphs depict mean ± SE. Box plots depict median (center line), mean (black box), quartiles, and 10-90% range (whiskers).

**Supplementary Figure 2. Optic fiber placements for fiber photometry experiments.** Optic fiber placements indicated by black dots for fiber photometry experiments during paired and unpaired fear conditioning described in Fig. 2 and Supplementary Fig. 4. Each subject was implanted with a single optic fiber (400 μm core diameter), but hemisphere was counterbalanced for equal representation.

**Supplementary Figure 3. SST-IN activity is not modulated by conditioned context-evoked freezing. a,** For $Ca^{2+}$-based imaging of SST-IN activity, SST-IRES-Cre mice (n = 6) received injections of conditional vector encoding GCamp6f and were implanted with a single optic fiber (400 μm core diameter) directed at PL. After surgical recovery, mice were subjected to unpaired fear conditioning, which entailed exposure in context A to 6 CS trials (2 KHz, 80 db, 20 s) and 6 US trials (0.7 mA, 2 s) in behavioral sessions separated by 15 min. At 24 and 48 hrs after conditioning, freezing was examined in response to a novel context (context B) or the original training arena (context A), respectively. **b**, Mean percent time freezing during pre-training baseline in context A, as well as post-conditioning exposures to contexts B and A, as function of time. Bars depict the average freezing for the 30-s period prior to the indicated time. **c**, Mean percent time freezing for duration of exposures depicted in (**b**). Effect of context: $F_{2,10} = 178.72$, $p = 1.49 \times 10^{-8}$, 1-way repeated measures ANOVA. **d**, US-related fluorescence signals during unpaired footshock trials in a representative animal. Scale = 30% ΔF/F x 2 s. **e**, Analysis of freezing-related fluorescence signals during exposure to contexts B and A. Freezing epochs were manually detected and registered to the entire fluorescence trace. Right: mean fluorescence trace for all aligned freezing epochs. **f**, Comparison of fluorescence signal changes associated with US exposure as well as aligned freezing epochs. All analyses were restricted to the initial 2 s after US or freezing initiation, and were based on normalization to the preceding 2 s of baseline. Effect of US versus freezing: $F_{2,10} = 59.33$, $p = 2.84 \times 10^{-6}$, 1-way repeated measures ANOVA. **g**, Mean frequency of fluorescence events (calcium transients) as function of context exposure. Effect of context: $F_{2,10} = 1.45$, $p = 0.29$, 1-way repeated measures ANOVA. *** $p < 0.001$ by Tukey's post-hoc test. Bar graphs depict mean ± SE.

**Supplementary Figure 4. CS-evoked freezing for cFos immunolabeling. a,** CS-evoked freezing for control subjects presented with 4 CSs without prior fear conditioning. **b,** CS-evoked freezing for control subjects that underwent auditory fear conditioning without subsequent presentation of 4 CSs. n = 4 mice. **c,** CS-evoked freezing for experimental subjects that were subjected to auditory fear conditioning followed, 24 hrs later, by presentation of 4 CSs. BL = baseline.

**Supplementary Figure 5. Fear learning does not lead to increased CS-evoked cFos expression in prelimbic PV-INs. a,** cFos labeling in SST-INs following CS exposure (4 trials, 20 s duration) at 24 hrs after fear conditioning (n = 3 mice) as compared to a control condition in which conditioning was omitted (n = 3 mice). **b,** CS-evoked freezing during the retrieval test for experimental animals and no conditioning controls. **c**, Proportion of $Tomato^{+}$ PV-INs that were co-labeled for cFos in each group. Total prelimbic (PL) counts, effect of training: $U = 7$, $p = 0.40$, Mann-Whitney *U* test. **d**, Percentage of PV-INs exhibiting cFos immunoreactivity as a function of behavioral training. **e**, Percentage of PV-INs exhibiting cFos immunoreactivity as a function of behavioral training, depicted as a pie slice (solid area).  Bar graphs depict mean ± SE.

**Supplementary Figure 6. Electrophysiological validation and optic fiber placement for SST-IN photoexcitation and inhibition. a,** Whole-cell recording from SST-INs expressing GFP-tagged Arch. n = 11 cells (3 mice). Graph and accompanying trace indicate that optic illumination (532 nm, constant, 20 s duration) hyperpolarized SST-INs and suppressed spontaneous firing. Scale = 20 mV x 5 s. **b,** Whole-cell recording from SST-INs expressing EYFP-tagged channelrhodopsin-2. n = 12 cells (3 mice). Example trace indicates that optic illumination (460 nm, 1 ms duration, 20 Hz) elicited reliable action potentials. Scale = 20 mV x 50 ms. Optic fiber placements for *in vivo* optogenetic manipulations of SST-INs are indicated for fear conditioned (**c**) and naïve (**d**) mice. Placements separately indicated for mice expressing Cre-dependent ChR2 (blue) and corresponding eYFP controls (black), as well as mice expressing Cre-dependent Arch (green) and corresponding eYFP controls (gray). Bar graphs depict mean ± SE.

**Supplementary Figure 7: No effect of SST-IN photoexcitation on locomotor activity in the open field test.** To assess potential locomotor effects of prelimbic SST-IN photoexcitation independent of fear conditioning, SST-IRES-Cre mice received prelimbic injections of conditional AAV-ChR2 or –eYFP control vectors, as in Fig. 4. Following viral incubation, mice were placed into an open field arena, and locomotor behavior was analyzed for 20 min, during which $light_{OFF}$ and $light_{ON}$ (473 nm, 10 ms pulse, 20 Hz) epochs lasting 5 min were counterbalanced to control for any ordering effects. **a**, Example activity plots for ChR2 (n = 5 mice) and eYFP controls (n = 5 mice) during $light_{OFF}$ and $light_{ON}$ epochs. **b-g**, No effect of photoexcitation on standard locomotor metrics in either ChR2 or eYFP subjects. Significance between $light_{OFF}$ and $light_{ON}$ epochs was established by paired t-test. **b**, Distance moved, effect of photoexcitation. ChR2: $t_4 = 0.097$, $p = 0.93$. eYFP: $t_4 = 0.42$, $p = 0.69$. **c**, Number of movement epochs, effect of photoexcitation. ChR2: $t_4 = 1.33$, $p = 0.25$. eYFP: $t_4 = 0.89$, p =

0.42. **d**, Number of rest epochs, effect of photoexcitation. ChR2: $t_4 = 1.25$, $p = 0.28$. eYFP: $t_4 = 0.71$, $p = 0.52$. **e**, Velocity during movement, effect of photoexcitation. ChR2: $t_4 = 0.35$, $p = 0.74$. eYFP: $t_4 = 0.45$, $p = 0.67$. **f**, Percent time moving, effect of photoexcitation. ChR2: $t_4 = 0.16$, $p = 0.88$. eYFP: $t_4 = 1.21$, $p = 0.29$. **g**, Percent time resting, effect of photoexcitation. ChR2: $t_4 = 0.16$, $p = 0.88$. eYFP: $t_4 = 1.22$, $p = 0.29$. Bar graphs depict mean ± SE.

**Supplementary Figure 8. Fiber placements and additional physiological data for SST-IN photoinhibition during learning. a**, Optic fiber placements for *in vivo* optogenetic manipulations of SST-INs, as described in Fig. 4a-c, are indicated for mice expressing Cre-dependent Arch (blue) and corresponding eYFP controls (black). **b**, For SST-INs, amplitude of sEPSCs depicted in Fig. 4d. Effect of virus: $t_{18} = 2.45$, $p = 0.025$, unpaired t-test; Arch, n = 10 cells (3 mice); eYFP, n= 10 cells (3 mice). * $p < 0.05$ by t-test. Bar graph depicts mean ± SE.

**Supplementary Figure 9. Fiber placements and additional physiological data for BLA photoinhibition during learning. a**, Optic fiber placements for *in vivo* optogenetic manipulations of BLA projection neurons, as described in Fig. 4f-h, are indicated for mice expressing Cre-dependent Arch (blue) and corresponding eYFP controls (black). **b**, For SST-INs, amplitude of sEPSCs depicted in Fig. 4i. Effect of virus: $t_{18} = 1.89$, $p = 0.070$, unpaired t-test; Arch, n = 14 cells (3 mice); eYFP, n= 13 cells (4 mice). Bar graph depicts mean ± SE.

**Supplementary Figure 10. CS-evoked freezing during learning for interrogation of synaptic connections between SST- and PV-INs.** Auditory fear conditioning entailed 6 presentations of CS and US in a paired or unpaired configuration. 24hrs after training, mice were sacrificed for brain slice electrophysiology, described in Fig. 6. **a**, Percent time freezing during CS trials is plotted for SST-FlpO/ PV-Cre/ Ai9 mice that were subjected to paired (n = 4 mice) or unpaired (n = 3 mice) auditory fear conditioning. **b**, Percent time freezing during CS trials is plotted for PV-FlpO/ SST-Cre/ Ai9 mice that were subjected to paired (n = 3 mice) or unpaired (n = 3 mice) auditory fear conditioning. BL = baseline.

**Supplementary Figure 11. Electrophysiological validation, viral expression and optic fiber placement for PV- and SST-IN photoexcitation and inhibition. a,** Whole-cell recording from PV-INs expressing GFP-tagged archaerhodopsin. n = 7 cells (3 mice). Graph and accompanying trace indicate that optic illumination (532 nm, constant, 20 s duration) hyperpolarized PV-INs and suppressed spike trains elicited by step depolarization. Scale = 20 mV x 5 s. (**b-c**) Viral expression (left) and optic fiber placements (right) related to *in vivo* optogenetic manipulations of PV- and SST-INs are indicated for conditioned PV-IRES-Cre (**b**) and SST-IRES-Cre x PV-IRES-Cre (**c**) mice. Placements separately indicated for ChR2-expressing mice (blue) and corresponding eYFP controls (black), as well as Arch-expressing mice (green) and corresponding eYFP controls (black). Scale = 500 μm. Bar graph depicts mean ± SE.

**Supplementary Figure 12. Comparison of compound excitatory and inhibitory postsynaptic currents elicited by BLA afferent stimulation in prelimbic cell types.** A CaMKII promoter-dependent ChR2 vector was injected into the BLA of PV- or SST-IRES-Cre mice crossed to the Ai9 reporter line. Whole-cell recordings were obtained from Tomato-positive PV- or SST-INs as well as surrounding PNs during photoexcitation of ChR2-expressing BLA afferents (460 nm, 1 ms pulse, 0.1 Hz). Excitatory and inhibitory postsynaptic currents were isolated by clamping at the reversal potential for GABA and glutamate receptors, respectively, in a low-chloride internal solution. **a,** Example raw (black) and mean traces (red) from each cell type, containing compound polysynaptic activity. Scale = 100 pA x 20 ms. **b,** Comparison of total charge transferred at each holding potential in SST-INs, n = 14 cells (4 mice), versus PNs, n = 12 cells (4 mice). EPSCs (-70 mV), effect of cell type: $t_{24} = 1.51$, $p = 0.14$, unpaired t-test.

IPSCs (0 mV), effect of cell type: *U* = 151, 6.25 x $10^{-4}$, Mann-Whitney *U* test. **c**, Within-cell ratio of charge transferred during EPSCs (-70 mV)/ IPSCs (0 mV) for SST-INs and PNs in SST-IRES-Cre/ Ai9 mice. Effect of cell type: *U* = 34, p = 0.020, Mann-Whitney *U* test. **d**, Comparison of total charge transferred at each holding potential in PV-INs, n = 11 cells (3 mice), versus PNs, n = 9 cells (3 mice). EPSCs (-70 mV), effect of cell type: *U* = 61.5, p = 0.38, Mann-Whitney *U* test. IPSCs (0 mV), effect of cell type: $t_{18}$ = 1.64, p = 0.12, unpaired t-test. **e**, Within-cell ratio of charge transferred during EPSCs (-70 mV)/ IPSCs (0 mV) for PV-INs and PNs in PV-IRES-Cre/ Ai9 mice. Effect of cell type: *U* = 40.5, p = 0.80, Mann-Whitney *U* test. * p < 0.05, *** p < 0.001 by Mann-Whitney *U* test. Box plots depict median (center line), mean (black box), quartiles, and 10-90% range (whiskers).

**Supplementary Figure 13. CS-evoked freezing during learning for interrogation of BLA transmission onto prelimbic cell types.** Auditory fear conditioning entailed 6 presentations of CS and US in a paired or unpaired configuration. 24hrs after training, mice were sacrificed for brain slice electrophysiology, described in Fig. 7. **a**, Percent time freezing during CS trials is plotted for SST-Cre/ Ai9 mice that were subjected to paired (n = 4 mice) or unpaired (n = 3 mice) auditory fear conditioning. **b**, Percent time freezing during CS trials is plotted for PV-FlpO/ SST-Cre/ Ai9 mice that were subjected to paired (n = 3 mice) or unpaired (n = 3 mice) auditory fear conditioning. BL = baseline.

**Supplementary Figure 14. Optic fiber placements, behavior and supplementary images for network cFos analysis in conditioned mice.** Data are supplementary to the analysis in Fig. 9a-e. **a**, Percent time freezing during baseline (BL) period and during each of 6 CS-US trials during fear conditioning for ChR2 mice (n = 5) and eYFP controls (n = 5). Optic fiber placements depicted in Supplementary Fig. 5. **b**, Percent time freezing during the baseline period and during optic stimulation (473 nm, 10 ms pulse, 20 Hz, 6 x 20 s epochs) trials at 24 hrs after fear conditioning. ChR2, effect of stimulation: $t_4 = 5.70$, $p = 0.0047$, paired t-test. eYFP, effect of stimulation: $t_4 = 0.65$, $p = 0.55$, paired t-test. ChR2 versus eYFP, change in freezing during optic stimulation: *U* = 0, $p = 0.0079$, Mann-Whitney *U* test. **c**, Optic fiber placements for ChR2 (blue) and eYFP mice (black) used for analysis of optogenetic cFos induction in conditioned mice. **d,** Example images of cFos immunofluorescence in the dentate gyrus of the dorsal hippocampus (DG), paraventricular thalamus (PVT), and intero (IMD), lateral (MDL), central (MDC) and medial divisions of the mediodorsal thalamus (MDM), for statistical analysis described in Fig. 9e. **e**, Example images of cFos immunofluorescence in the caudate putamen (CPu), accumbens core (AcC), accumbens shell (AcS), and ventral hippocampus cornu ammonis area 1 (CA1), for statistical analysis described in Fig. 9e. Scale = 500 $\mu$m (CPu/ NAc images), 200 $\mu$m (all other images). ** $p < 0.01$ by paired t-test (**b**, left) or Mann-Whitney *U* test (**b**, right). Bar graphs depict mean ± SE. Box plots depict median (center line), quartiles, and 10-90% range (whiskers).

**Supplementary Figure 15. Optic fiber placements, behavior and supplementary images for network cFos analysis in naïve mice.** Data are supplementary to the analysis in Fig. 9f-h. **a**, Percent time freezing during the baseline period and during optic stimulation (473 nm, 10 ms pulse, 20 Hz, 6 x 20 s epochs) trials in naïve subjects expressing ChR2 (n = 5) or eYFP vectors (n = 5). ChR2, effect of stimulation: $t_4 = 2.02$, $p = 0.11$, paired t-test. eYFP, effect of stimulation: $t_4 = 0.91$, $p = 0.42$, paired t-test. ChR2 versus eYFP, change in freezing during optic stimulation: *U* = 17, $p = 0.42$, Mann-Whitney *U* test. **b**, Example images of cFos immunofluorescence from stimulated prelimbic cortex tissue in ChR2 mice (right) or eYFP control mice (left). Lower panels depict cells co-labeled for eYFP (green) and cFos (red). Scale = 500 $\mu$m (upper), 50 $\mu$m (lower). **c**, Example images of cFos immunofluorescence from stimulated animals for statistical analysis in Fig. 9h. LA = lateral, BA = basal amygdala. LHb = lateral habenula. MHb = medial habenula. PVT = paraventricular thalamus. DM = dorsomedial hypothalamus. CPu = caudate putamen.

NAcC = nucleus accumbens core. NAcS = nucleus accumbens shell. CA1 = cornu ammonis area 1 of the ventral hippocampus (CA1). DG = dentate gyrus of the dorsal hippocampus. MDM = medial mediodorsal thalamus. MDC = central mediodorsal thalamus. MDL = lateral mediodorsal thalamus. IMD = interomedial dorsal thalamus. Scale = 500 μm (CPu/ NAc images), 200 μm (all other images). Bar graphs depict mean ± SE. Box plots depict median (center line), quartiles, and 10-90% range (whiskers).

**Supplementary Fig. 16. Behavior and supplementary images for network cFos analysis following memory retrieval.** Data are supplementary to the analysis in Fig. 9i-j. **a**, Percent time freezing during CS-US pairing for conditioned (n = 6) and non-conditioned mice (n = 5). **b**, Percent time freezing during CS retrieval test. Conditioned mice, effect of CS: $t_5 = 8.23$, p = 0.00043, paired t-test. Non-conditioned mice, effect of CS: *W* = 0, p = 0.50, Wilcoxon rank sum test. Conditioned versus non-conditioned mice, change in freezing during CS exposure: *U* = 0, p = 0.0043. **c**, Example images of cFos immunofluorescence following CS exposure in conditioned (Cond) and non-conditioned (NC) mice, for statistical analysis in Fig. 9j. LA = lateral, BA = basal amygdala. LHb = lateral habenula. MHb = medial habenula. PVT = paraventricular thalamus. DM = dorsomedial hypothalamus. CA1 = cornu ammonis area 1 of the ventral hippocampus (CA1). **d**, Additional example images of cFos immunofluorescence following CS exposure in conditioned (Cond) and non-conditioned (NC) mice. CPu = caudate putamen. NAcC = nucleus accumbens core. NAcS = nucleus accumbens shell. DG = dentate gyrus of the dorsal hippocampus. MDM = medial mediodorsal thalamus. MDC = central mediodorsal thalamus. MDL = lateral mediodorsal thalamus. IMD = interomedial dorsal thalamus. Scale = 500 μm (CPu/ NAc images), 200 μm (all other images). **p < 0.01 by paired t-test (**b**, left) or Mann-Whitney *U* test (**b**, right). Bar graphs depict mean ± SE. Box plots depict median (center line), quartiles, and 10-90% range (whiskers).

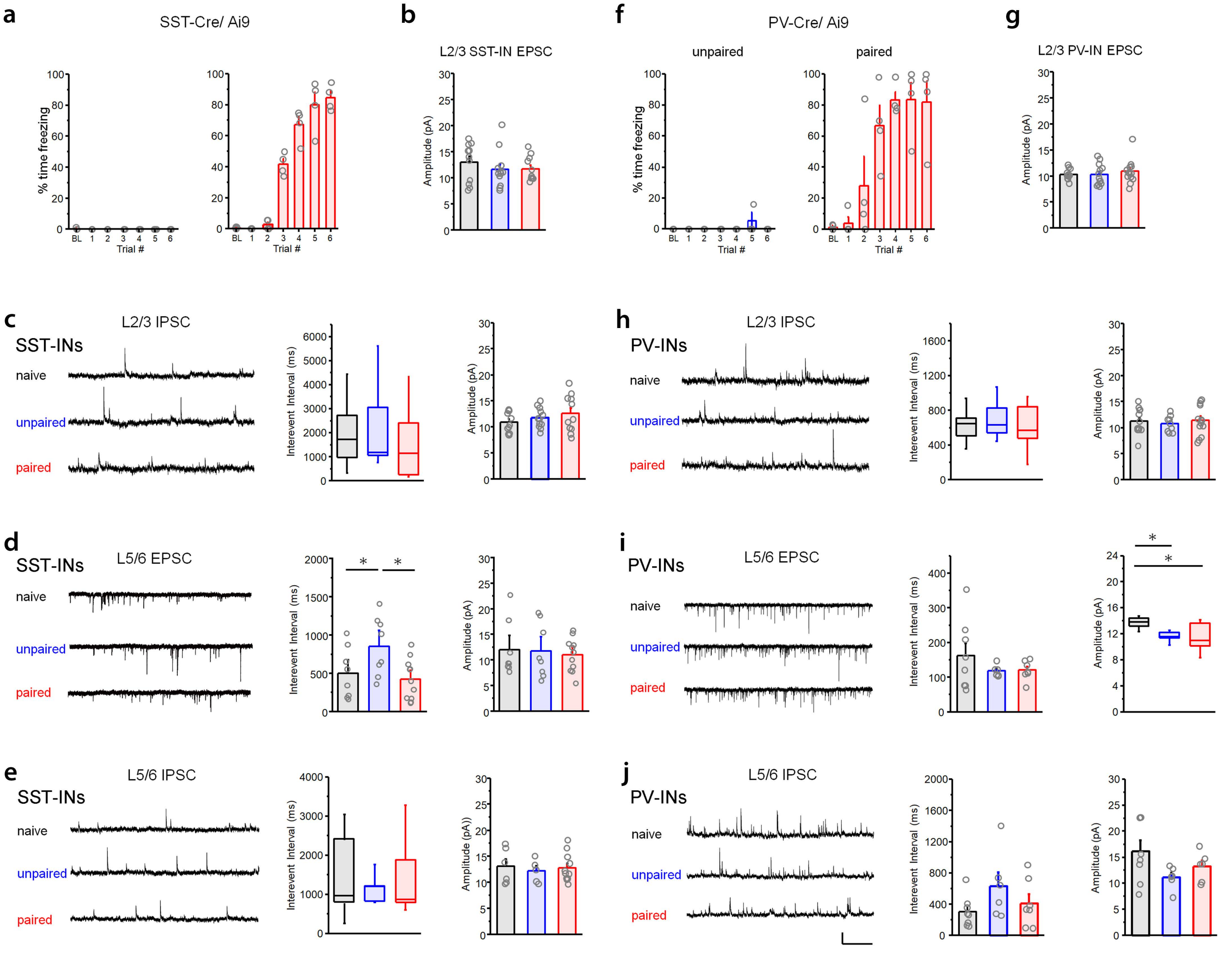


**Supplementary Figure 1. Spontaneous synaptic transmission in prelimbic interneurons after fear conditioning.** Example raw traces, mean interevent interval (IEI) and amplitude of events are shown. Whole-cell recordings were obtained from acute brain slices from naïve mice or littermates that were subjected paired or unpaired auditory fear conditioning, as in Fig. 1. Scale for all traces = 20 pA x 1 s. **a,** CS-evoked freezing during fear conditioning for SST-IRES-Cre/ Ai9 mice used for spontaneous transmission analysis. **b**, L2/3 SST-IN sEPSC amplitude, effect of training: $F_{2,31} = 0.58$, p = 0.56, 1-way ANOVA; naïve, n = 11 cells (3 mice); unpaired, n =12 cells (3 mice); paired, n = 11 cells (4 mice). **c**, L2/3 SST-IN sIPSCs. Interevent interval, effect of training: $\chi^2 = 1.35$ (2), p = 0.51, Kruskal-Wallis ANOVA; sIPSC amplitude, effect of training: $F_{2,31} = 1.18$, p = 0.32, 1-way ANOVA; naïve, n = 11 cells (3 mice); unpaired, n =12 cells (3 mice); paired, n = 11 cells (4 mice). **d**, L5/6 SST-IN sEPSCs. Interevent interval, effect of training: $F_{2,24} = 4.45$, p = 0.023, 1-way ANOVA; amplitude: $F_{2,24} = 0.12$, p = 0.89, 1-way ANOVA; naïve, n = 8 cells (3 mice); unpaired, n = 8 cells (3 mice); paired, n = 11 cells (3 mice). **e**, L5/6 SST-IN sIPSCs. Interevent interval, effect of training: $F_{2,19} = 0.12$, p = 0.89, 1-way ANOVA; amplitude, effect of training: $F_{2,19} = 0.15$, p = 0.86, 1-way ANOVA; naïve, n = 6 cells (3 mice); unpaired, n = 5 cells (3 mice); paired, n = 11 cells (3 mice). **f**, CS-evoked freezing during fear conditioning for PV-IRES-Cre/ Ai9 mice used for spontaneous transmission analysis. **g**, L2/3 PV-IN sEPSC amplitude, effect of training: $F_{2,36} = 0.45$, p = 0.64, 1-way ANOVA; naïve, n = 13 cells (3 mice); unpaired, n =13 cells (3 mice); paired, n = 13 cells (3 mice). **h**, L2/3 PV-IN sIPSCs. Interevent interval, effect of training: $F_{2,31} = 0.23$, p = 0.80, 1-way ANOVA; amplitude: $F_{2,31} = 0.22$, p = 0.80, 1-way ANOVA; naïve, n = 12 cells (3 mice); unpaired, n = 9 cells (3 mice); paired, n = 13 cells (3 mice). **i**, L5/6 PV-IN sEPSCs. Interevent interval, effect of training: $F_{2,18} = 1.04$, p = 0.37, 1-way ANOVA; amplitude, effect of training: $F_{2,18} = 5.41$, p = 0.014, 1-way ANOVA; naïve, n = 8 cells (3 animals); unpaired, n = 6 cells (3 animals); paired, n = 7 cells (3 animals). **j**, L5/6 PV-IN sIPSCs. Interevent interval, effect of training: $F_{2,18} = 1.83$, p = 0.19, 1-way ANOVA; amplitude, effect of training: $F_{2,18} = 2.48$, p = 0.11, 1-way ANOVA; naïve, n = 8 cells (3 animals); unpaired, n = 6 cells (3 animals); n = 7 cells (3 animals). * $p < 0.05$ by Tukey's post-hoc test (**d**) or Dunn's post-hoc test (**i**). Bar graphs depict mean ± SE. Box plots depict median (center line), mean (black box), quartiles, and 10-90% range (whiskers).

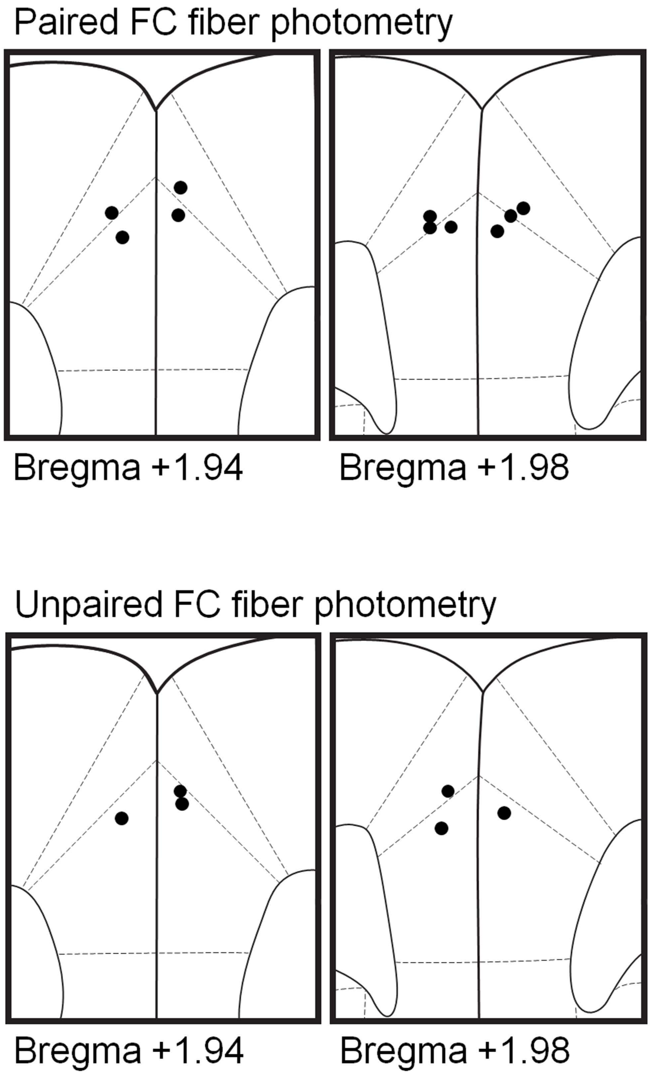


**Supplementary Figure 2. Optic fiber placements for fiber photometry experiments.** Optic fiber placements indicated by black dots for fiber photometry experiments during paired and unpaired fear conditioning described in Fig. 2 and Supplementary Fig. 4. Each subject was implanted with a single optic fiber (400 μm core diameter), but hemisphere was counterbalanced for equal representation.

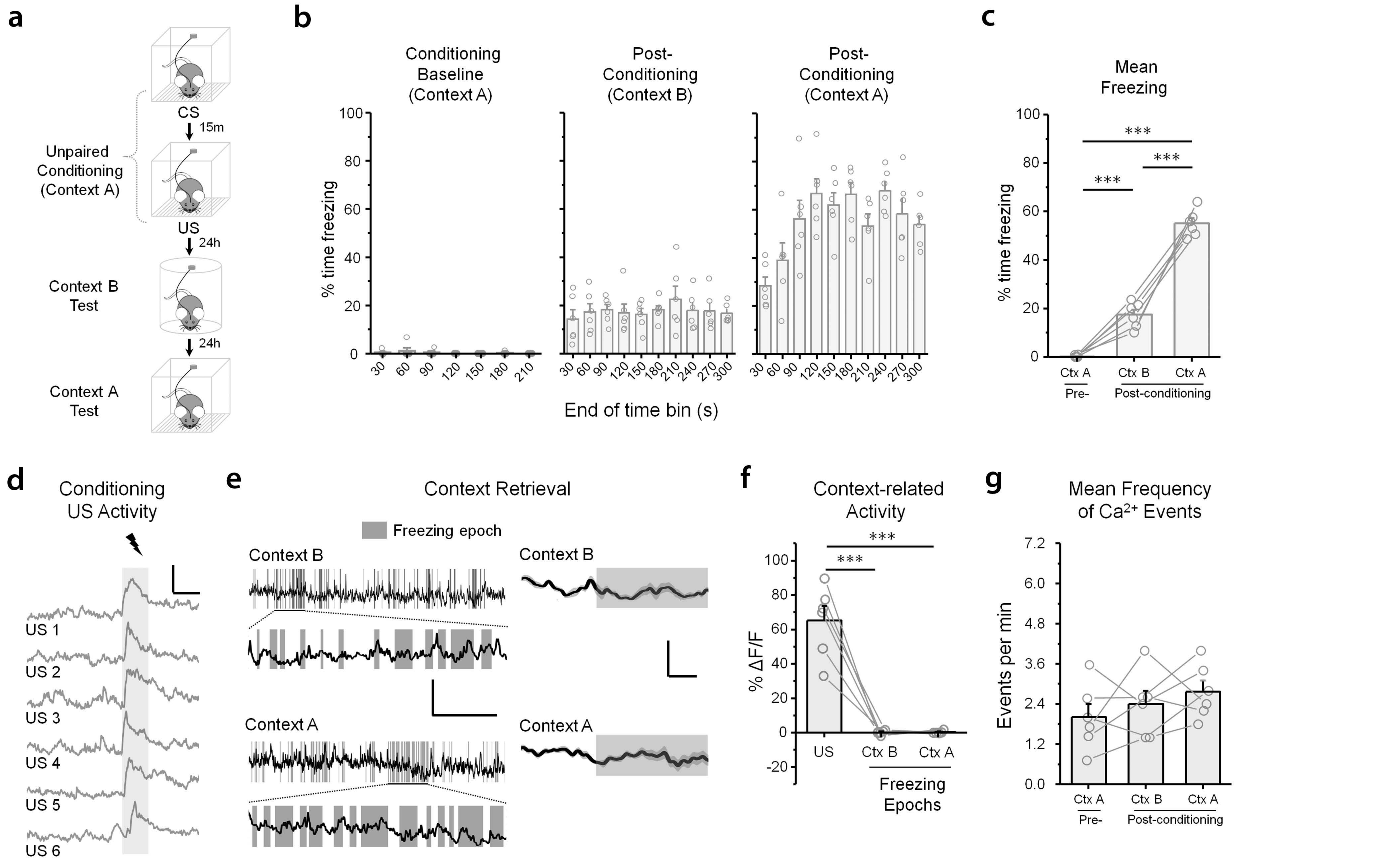


**Supplementary Figure 3. SST-IN activity is not modulated by conditioned context-evoked freezing. a,** For $Ca^{2+}$-based imaging of SST-IN activity, SST-IRES-Cre mice (n = 6) received injections of conditional vector encoding GCamp6f and were implanted with a single optic fiber (400 $\mu$m core diameter) directed at PL. After surgical recovery, mice were subjected to unpaired fear conditioning, which entailed exposure in context A to 6 CS trials (2 KHz, 80 db, 20 s) and 6 US trials (0.7 mA, 2 s) in behavioral sessions separated by 15 min. At 24 and 48 hrs after conditioning, freezing was examined in response to a novel context (context B) or the original training arena (context A), respectively. **b**, Mean percent time freezing during pre-training baseline in context A, as well as post-conditioning exposures to contexts B and A, as function of time. Bars depict the average freezing for the 30-s period prior to the indicated time. **c**, Mean percent time freezing for duration of exposures depicted in (**b**). Effect of context: $F_{2,10}$ = 178.72, p = 1.49 x $10^{-8}$, 1-way repeated measures ANOVA. **d**, US-related fluorescence signals during unpaired footshock trials in a representative animal. Scale = 30% ΔF/F x 2 s. **e**, Analysis of freezing-related fluorescence signals during exposure to contexts B and A. Freezing epochs were manually detected and registered to the entire fluorescence trace. Right: mean fluorescence trace for all aligned freezing epochs. **f**, Comparison of fluorescence signal changes associated with US exposure as well as aligned freezing epochs. All analyses were restricted to the initial 2 s after US or freezing initiation, and were based on normalization to the preceding 2 s of baseline. Effect of US versus freezing: $F_{2,10}$ = 59.33, p = 2.84 x $10^{-6}$, 1-way repeated measures ANOVA. **g**, Mean frequency of fluorescence events (calcium transients) as function of context exposure. Effect of context: $F_{2,10}$ = 1.45, p = 0.29, 1-way repeated measures ANOVA. *** p < 0.001 by Tukey's post-hoc test. Bar graphs depict mean ± SE.

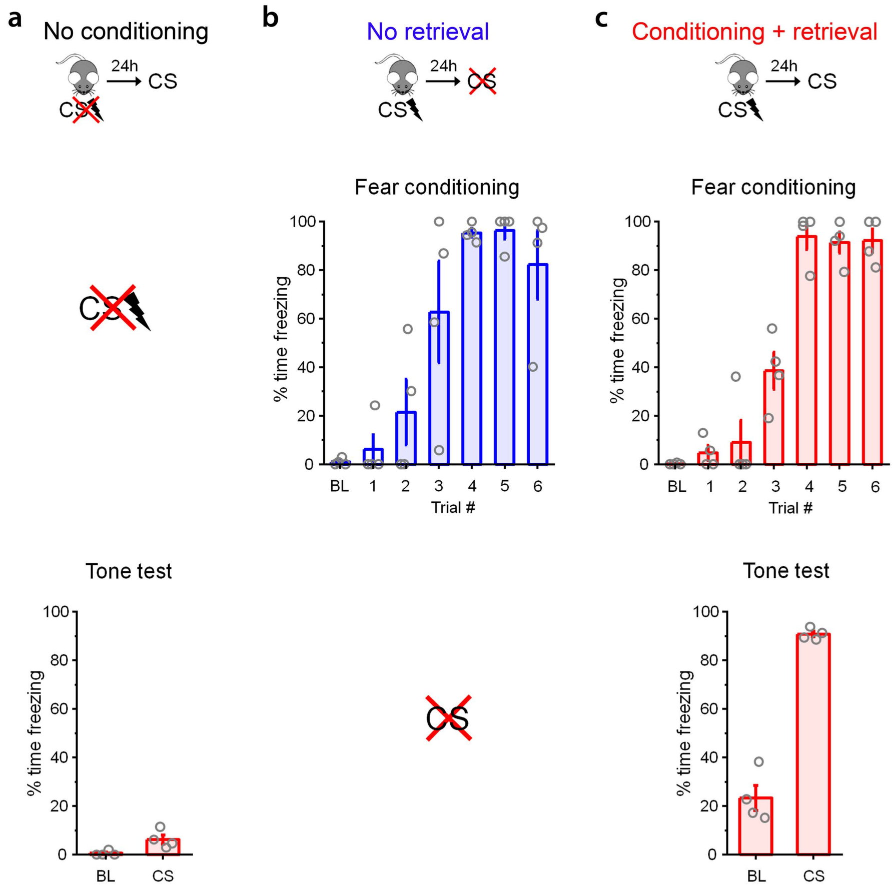


**Supplementary Figure 4. CS-evoked freezing for cFos immunolabeling. a,** CS-evoked freezing for control subjects presented with 4 CSs without prior fear conditioning. **b,** CS-evoked freezing for control subjects that underwent auditory fear conditioning without subsequent presentation of 4 CSs. n = 4 mice. **c,** CS-evoked freezing for experimental subjects that were subjected to auditory fear conditioning followed, 24 hrs later, by presentation of 4 CSs. BL = baseline.

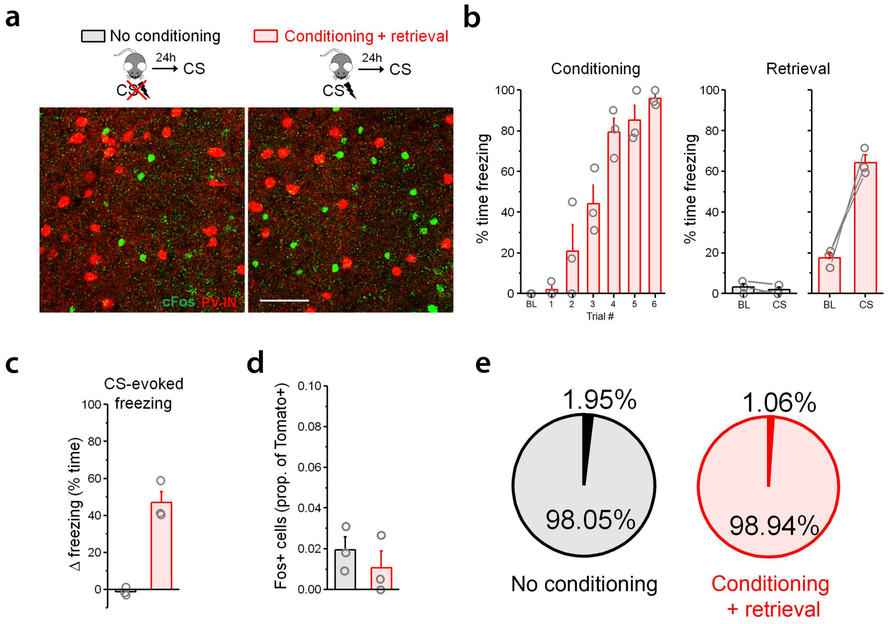


**Supplementary Figure 5. Fear learning does not lead to increased CS-evoked cFos expression in prelimbic PV-INs. a,** cFos labeling in SST-INs following CS exposure (4 trials, 20 s duration) at 24 hrs after fear conditioning (n = 3 mice) as compared to a control condition in which conditioning was omitted (n = 3 mice). **b,** CS-evoked freezing during the retrieval test for experimental animals and no conditioning controls. **c**, Proportion of Tomato$^+$ PV-INs that were co-labeled for cFos in each group. Total prelimbic (PL) counts, effect of training: $U = 7$, p = 0.40, Mann-Whitney *U* test. **d**, Percentage of PV-INs exhibiting cFos immunoreactivity as a function of behavioral training. **e**, Percentage of PV-INs exhibiting cFos immunoreactivity as a function of behavioral training, depicted as a pie slice (dark area). Bar graphs depict mean ± SE.

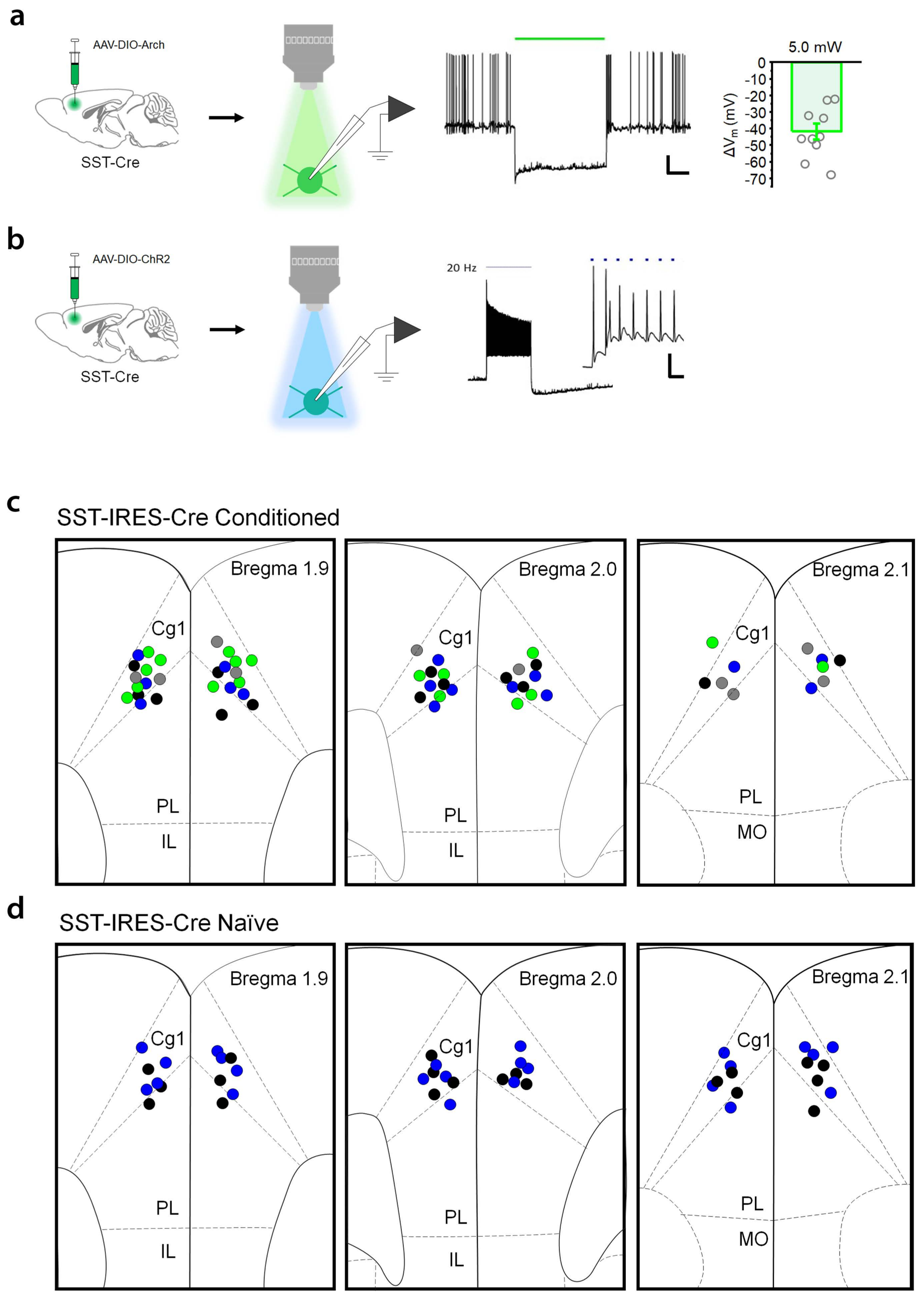


**Supplementary Figure 6. Electrophysiological validation and optic fiber placement for SST-IN photoexcitation and inhibition. a,** Whole-cell recording from SST-INs expressing GFP-tagged Arch. n = 11 cells (3 mice). Graph and accompanying trace indicate that optic illumination (532 nm, constant, 20 s duration) hyperpolarized SST-INs and suppressed spontaneous firing. Scale = 20 mV x 5 s. **b,** Whole-cell recording from SST-INs expressing EYFP-tagged channelrhodopsin-2. n = 12 cells (3 mice). Example trace indicates that optic illumination (460 nm, 1 ms duration, 20 Hz) elicited reliable action potentials. Scale = 20 mV x 50 ms. Optic fiber placements for *in vivo* optogenetic manipulations of SST-INs are indicated for fear conditioned (**c**) and naïve (**d**) mice. Placements separately indicated for mice expressing Cre-dependent ChR2 (blue) and corresponding eYFP controls (black), as well as mice expressing Cre-dependent Arch (green) and corresponding eYFP controls (gray). Bar graphs depict mean ± SE.

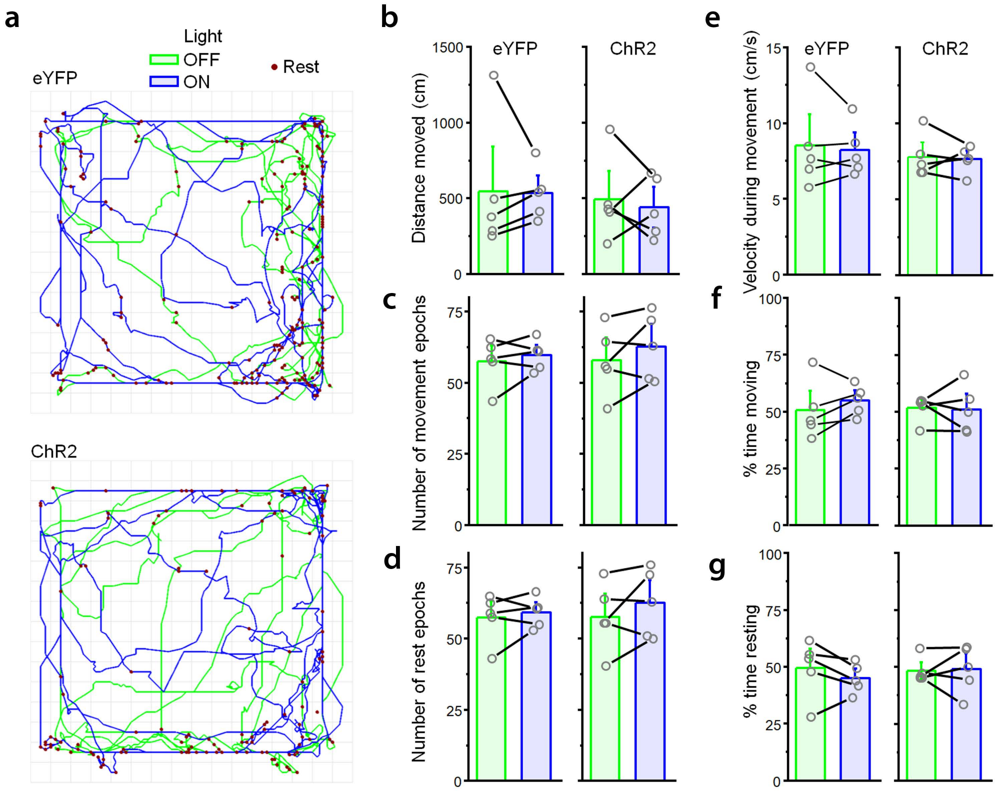


**Supplementary Figure 7: No effect of SST-IN photoexcitation on locomotor activity in the open field test.** To assess potential locomotor effects of prelimbic SST-IN photoexcitation independent of fear conditioning, SST-IRES-Cre mice received prelimbic injections of conditional AAV-ChR2 or –eYFP control vectors, as in Fig. 4. Following viral incubation, mice were placed into an open field arena, and locomotor behavior was analyzed for 20 min, during which $light_{OFF}$ and $light_{ON}$ (473 nm, 10 ms pulse, 20 Hz) epochs lasting 5 min were counterbalanced to control for any ordering effects. **a**, Example activity plots for ChR2 (n = 5 mice) and eYFP controls (n = 5 mice) during $light_{OFF}$ and $light_{ON}$ epochs. **b-g**, No effect of photoexcitation on standard locomotor metrics in either ChR2 or eYFP subjects. Significance between $light_{OFF}$ and $light_{ON}$ epochs was established by paired t-test. **b**, Distance moved, effect of photoexcitation. ChR2: $t_4 = 0.097$, $p = 0.93$. eYFP: $t_4 = 0.42$, $p = 0.69$. **c**, Number of movement epochs, effect of photoexcitation. ChR2: $t_4 = 1.33$, $p = 0.25$. eYFP: $t_4 = 0.89$, $p = 0.42$. **d**, Number of rest epochs, effect of photoexcitation. ChR2: $t_4 = 1.25$, $p = 0.28$. eYFP: $t_4 = 0.71$, $p = 0.52$. **e**, Velocity during movement, effect of photoexcitation. ChR2: $t_4 = 0.35$, $p = 0.74$. eYFP: $t_4 = 0.45$, $p = 0.67$. **f**, Percent time moving, effect of photoexcitation. ChR2: $t_4 = 0.16$, $p = 0.88$. eYFP: $t_4 = 1.21$, $p = 0.29$. **g**, Percent time resting, effect of photoexcitation. ChR2: $t_4 = 0.16$, $p = 0.88$. eYFP: $t_4 = 1.22$, $p = 0.29$. Bar graphs depict mean ± SE.

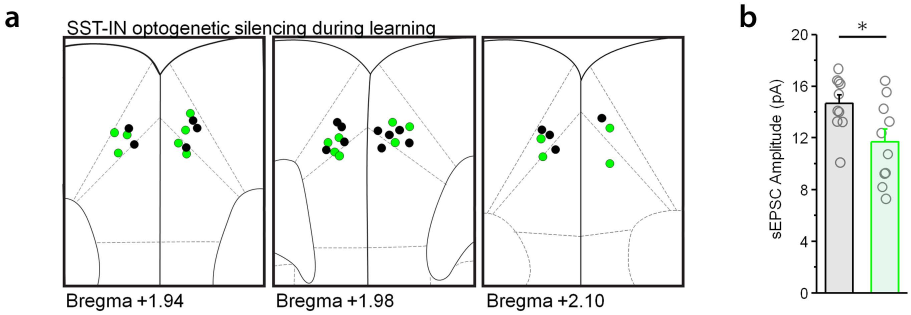


**Supplementary Figure 8. Fiber placements and additional physiological data for SST-IN photoinhibition during learning. a**, Optic fiber placements for *in vivo* optogenetic manipulations of SST-INs, as described in Fig. 4a-c, are indicated for mice expressing Cre-dependent Arch (blue) and corresponding eYFP controls (black). **b**, For SST-INs, amplitude of sEPSCs depicted in Fig. 4d. Effect of virus: $t_{18} = 2.45$, $p = 0.025$, unpaired t-test; Arch, n = 10 cells (3 mice); eYFP, n= 10 cells (3 mice). * $p < 0.05$ by t-test. Bar graph depicts mean ± SE.

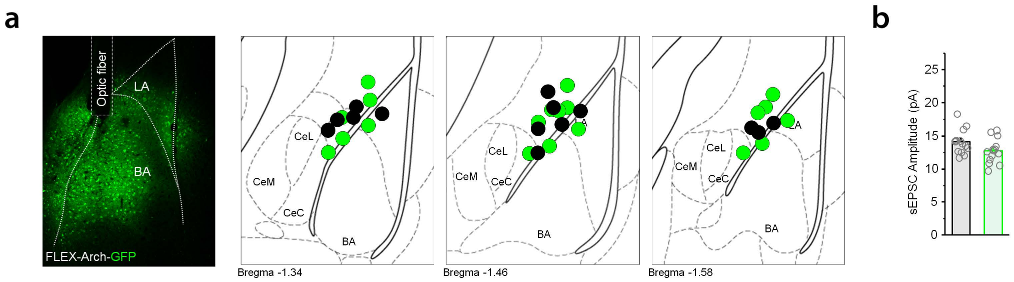


**Supplementary Figure 9. Fiber placements and additional physiological data for BLA photoinhibition during learning. a**, Optic fiber placements for *in vivo* optogenetic manipulations of BLA projection neurons, as described in Fig. 4f-h, are indicated for mice expressing Cre-dependent Arch (blue) and corresponding eYFP controls (black). **b**, For SST-INs, amplitude of sEPSCs depicted in Fig. 4i. Effect of virus: $t_{18}$ = 1.89, p = 0.070, unpaired t-test; Arch, n = 14 cells (3 mice); eYFP, n= 13 cells (4 mice). Bar graph depicts mean ± SE.

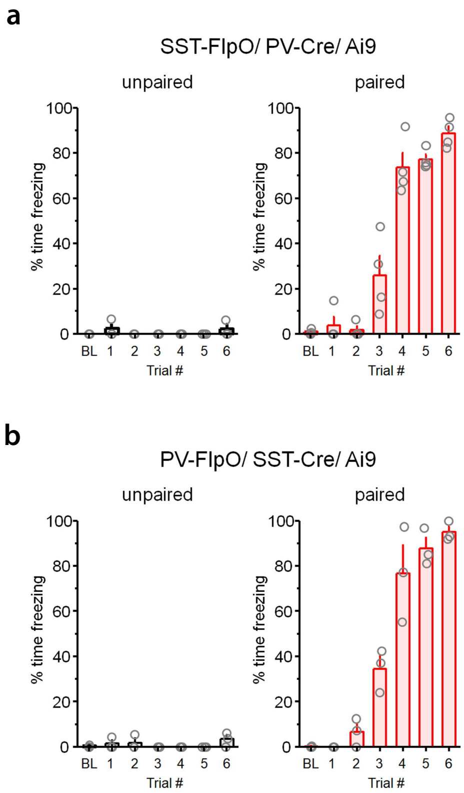


**Supplementary Figure 10. CS-evoked freezing during learning for interrogation of synaptic connections between SST- and PV-INs.** Auditory fear conditioning entailed 6 presentations of CS and US in a paired or unpaired configuration. 24hrs after training, mice were sacrificed for brain slice electrophysiology, described in Fig. 6. **a**, Percent time freezing during CS trials is plotted for SST-FlpO/ PV-Cre/ Ai9 mice that were subjected to paired (n = 4 mice) or unpaired (n = 3 mice) auditory fear conditioning. **b**, Percent time freezing during CS trials is plotted for PV-FlpO/ SST-Cre/ Ai9 mice that were subjected to paired (n = 3 mice) or unpaired (n = 3 mice) auditory fear conditioning. BL = baseline.

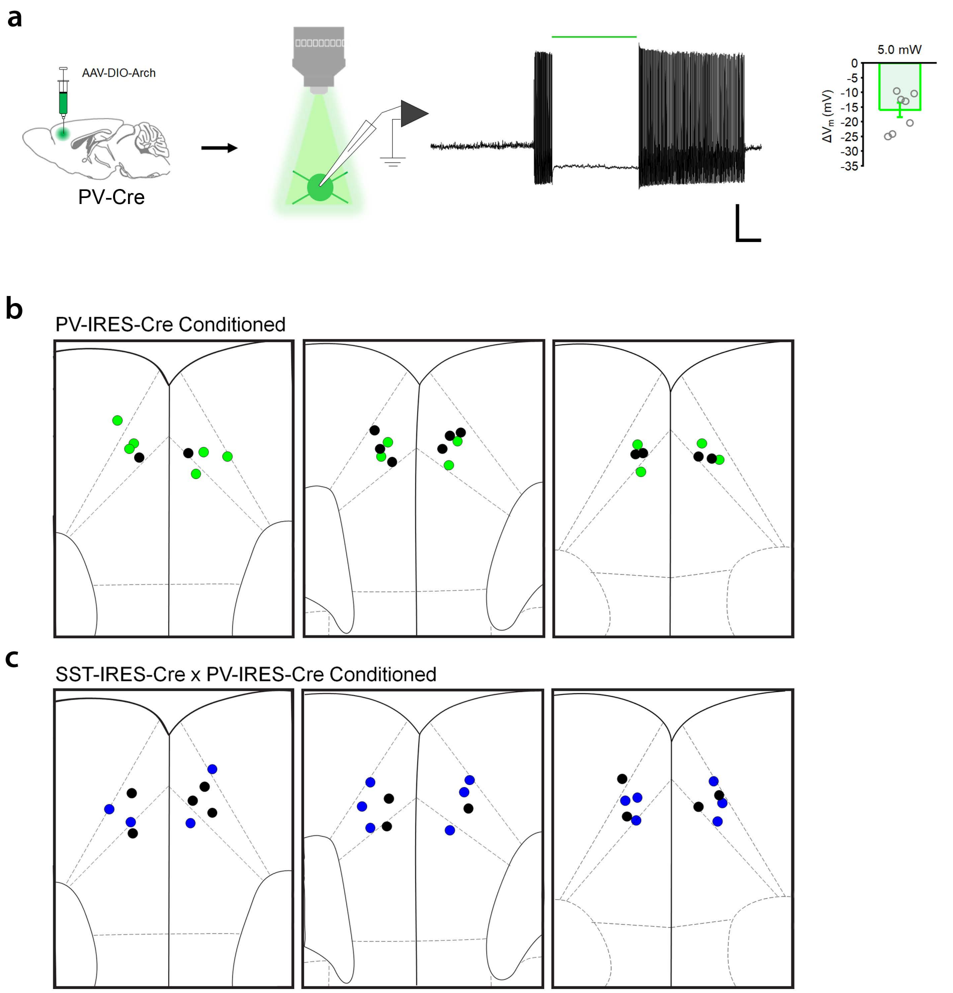


**Supplementary Figure 11. Electrophysiological validation, viral expression and optic fiber placement for PV- and SST-IN photoexcitation and inhibition. a,** Whole-cell recording from PV-INs expressing GFP-tagged archaerhodopsin. n = 7 cells (3 mice). Graph and accompanying trace indicate that optic illumination (532 nm, constant, 20 s duration) hyperpolarized PV-INs and suppressed spike trains elicited by step depolarization. Scale = 20 mV x 5 s. (**b-c**) Viral expression (left) and optic fiber placements (right) related to *in vivo* optogenetic manipulations of PV- and SST-INs are indicated for conditioned PV-IRES-Cre (**b**) and SST-IRES-Cre x PV-IRES-Cre (**c**) mice. Placements separately indicated for ChR2-expressing mice (blue) and corresponding eYFP controls (black), as well as Arch-expressing mice (green) and corresponding eYFP controls (black). Scale = 500 μm. Bar graph depicts mean ± SE.

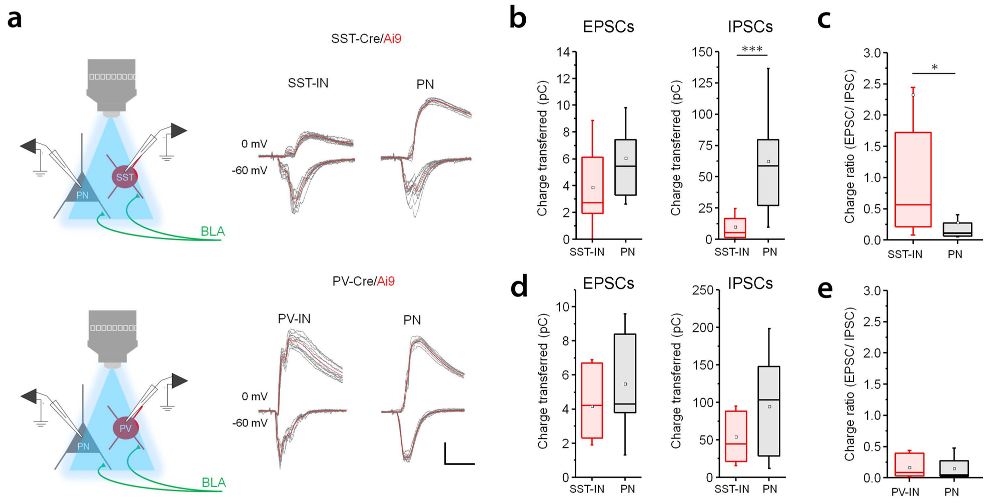


**Supplementary Figure 12. Comparison of compound excitatory and inhibitory postsynaptic currents elicited by BLA afferent stimulation in prelimbic cell types.** A CaMKII promoter-dependent ChR2 vector was injected into the BLA of PV- or SST-IRES-Cre mice crossed to the Ai9 reporter line. Whole-cell recordings were obtained from Tomato-positive PV- or SST-INs as well as surrounding PNs during photoexcitation of ChR2-expressing BLA afferents (460 nm, 1 ms pulse, 0.1 Hz). Excitatory and inhibitory postsynaptic currents were isolated by clamping at the reversal potential for GABA and glutamate receptors, respectively, in a low-chloride internal solution. **a,** Example raw (black) and mean traces (red) from each cell type, containing compound polysynaptic activity. Scale = 100 pA x 20 ms. **b,** Comparison of total charge transferred at each holding potential in SST-INs, n = 14 cells (4 mice), versus PNs, n = 12 cells (4 mice). EPSCs (-70 mV), effect of cell type: $t_{24} = 1.51$, p = 0.14, unpaired t-test. IPSCs (0 mV), effect of cell type: $U$ = 151, 6.25 x $10^{-4}$, Mann-Whitney *U* test. **c**, Within-cell ratio of charge transferred during EPSCs (-70 mV)/ IPSCs (0 mV) for SST-INs and PNs in SST-IRES-Cre/ Ai9 mice. Effect of cell type: $U$ = 34, p = 0.020, Mann-Whitney *U* test. **d**, Comparison of total charge transferred at each holding potential in PV-INs, n = 11 cells (3 mice), versus PNs, n = 9 cells (3 mice). EPSCs (-70 mV), effect of cell type: $U$ = 61.5, p = 0.38, Mann-Whitney *U* test. IPSCs (0 mV), effect of cell type: $t_{18} = 1.64$, p = 0.12, unpaired t-test. **e**, Within-cell ratio of charge transferred during EPSCs (-70 mV)/ IPSCs (0 mV) for PV-INs and PNs in PV-IRES-Cre/ Ai9 mice. Effect of cell type: $U$ = 40.5, p = 0.80, Mann-Whitney *U* test. * $p < 0.05$, *** $p < 0.001$ by Mann-Whitney *U* test. Box plots depict median (center line), mean (black box), quartiles, and 10-90% range (whiskers).

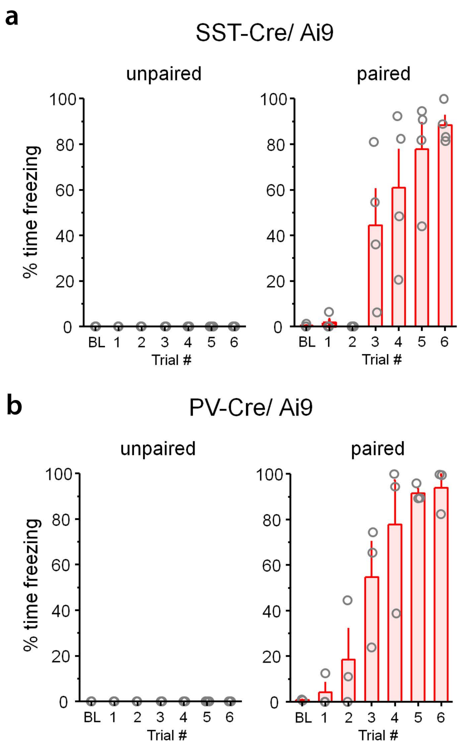


**Supplementary Figure 13. CS-evoked freezing during learning for interrogation of BLA transmission onto prelimbic cell types.** Auditory fear conditioning entailed 6 presentations of CS and US in a paired or unpaired configuration. 24hrs after training, mice were sacrificed for brain slice electrophysiology, described in Fig. 7. **a**, Percent time freezing during CS trials is plotted for SST-Cre/ Ai9 mice that were subjected to paired (n = 4 mice) or unpaired (n = 3 mice) auditory fear conditioning. **b**, Percent time freezing during CS trials is plotted for PV-FlpO/ SST-Cre/ Ai9 mice that were subjected to paired (n = 3 mice) or unpaired (n = 3 mice) auditory fear conditioning. BL = baseline.

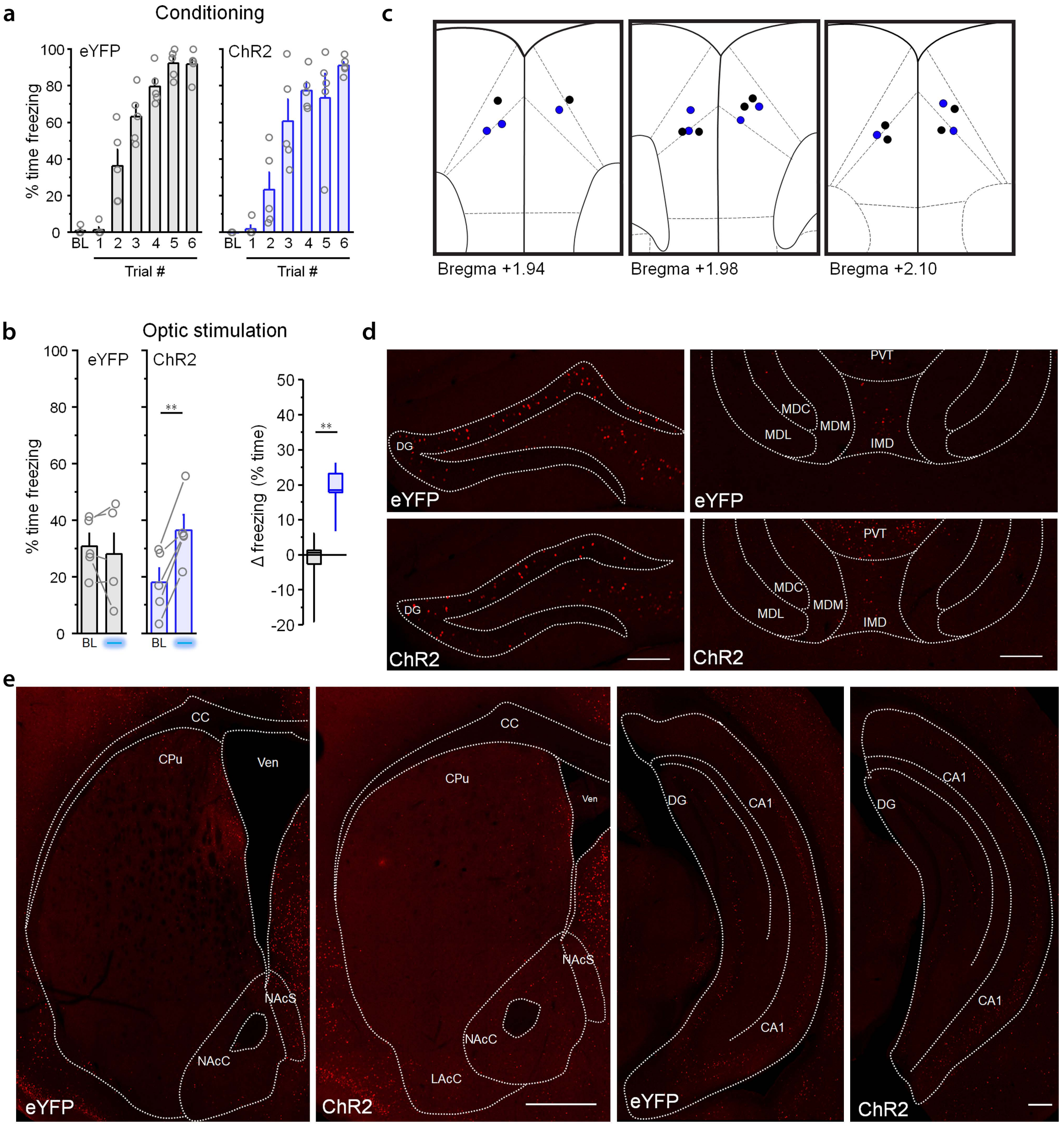


**Supplementary Figure 14. Optic fiber placements, behavior and supplementary images for network cFos analysis in conditioned mice.** Data are supplementary to the analysis in Fig. 9a-e. **a**, Percent time freezing during baseline (BL) period and during each of 6 CS-US trials during fear conditioning for ChR2 mice (n = 5) and eYFP controls (n = 5). Optic fiber placements depicted in Supplementary Fig. 5. **b**, Percent time freezing during the baseline period and during optic stimulation (473 nm, 10 ms pulse, 20 Hz, 6 x 20 s epochs) trials at 24 hrs after fear conditioning. ChR2, effect of stimulation: $t_4 = 5.70$, $p = 0.0047$, paired t-test. eYFP, effect of stimulation: $t_4 = 0.65$, $p = 0.55$, paired t-test. ChR2 versus eYFP, change in freezing during optic stimulation: $U = 0$, $p = 0.0079$, Mann-Whitney *U* test. **c**, Optic fiber placements for ChR2 (blue) and eYFP mice (black) used for analysis of optogenetic cFos induction in conditioned mice. **d,** Example images of cFos immunofluorescence in the dentate gyrus of the dorsal hippocampus (DG), paraventricular thalamus (PVT), and intero (IMD), lateral (MDL), central (MDC) and medial divisions of the mediodorsal thalamus (MDM), for statistical analysis described in Fig. 9e. **e**, Example images of cFos immunofluorescence in the caudate putamen (CPu), accumbens core (AcC), accumbens shell (AcS), and ventral hippocampus cornu ammonis area 1 (CA1), for statistical analysis described in Fig. 9e. Scale = 500 μm (CPu/ NAc images), 200 μm (all other images). ** $p < 0.01$ by paired t-test (**b**, left) or Mann-Whitney *U* test (**b**, right). Bar graphs depict mean ± SE. Box plots depict median (center line), quartiles, and 10-90% range (whiskers).

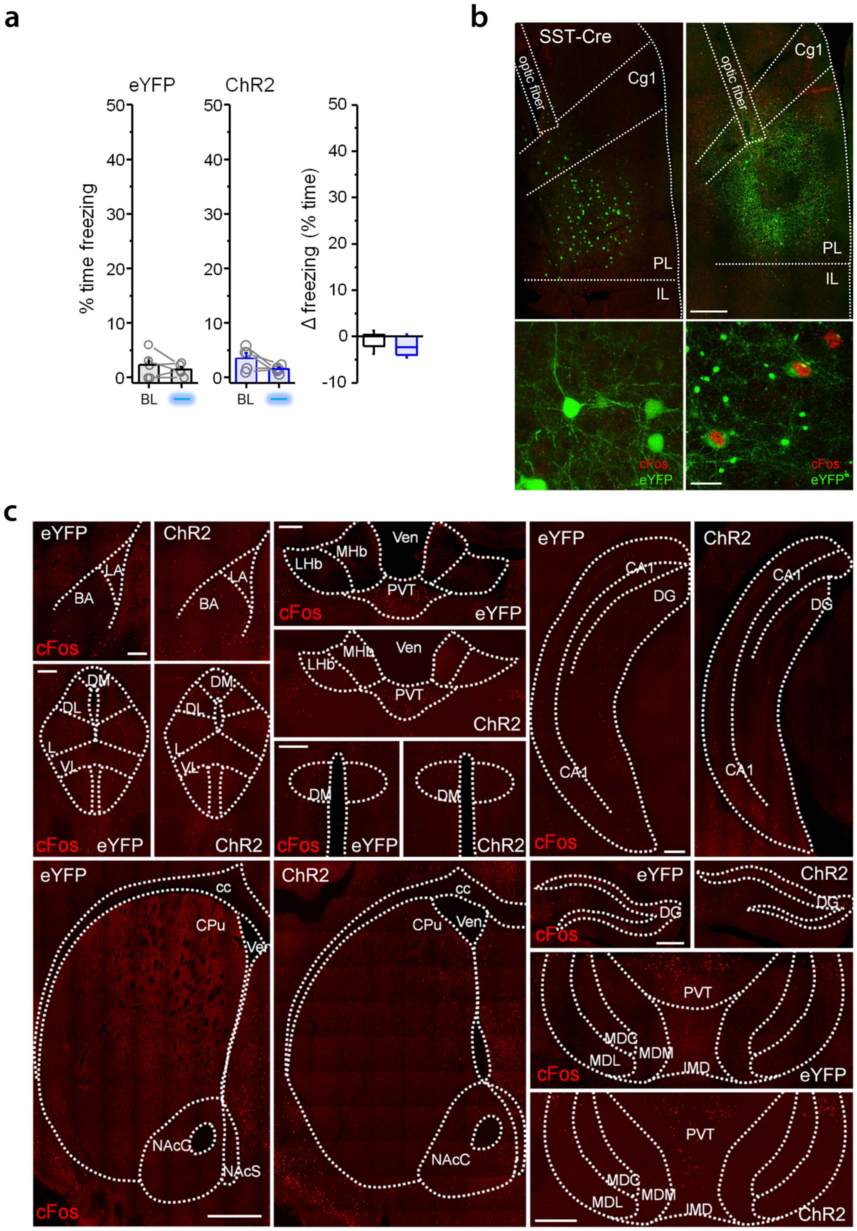

**Supplementary Figure 15. Optic fiber placements, behavior and supplementary images for network cFos analysis in naïve mice.** Data are supplementary to the analysis in Fig. 9f-h. **a**, Percent time freezing during the baseline period and during optic stimulation (473 nm, 10 ms pulse, 20 Hz, 6 x 20 s epochs) trials in naïve subjects expressing ChR2 (n = 5) or eYFP vectors (n = 5). ChR2, effect of stimulation: $t_4 = 2.02$, p = 0.11, paired t-test. eYFP, effect of stimulation: $t_4 = 0.91$, p = 0.42, paired t-test. ChR2 versus eYFP, change in freezing during optic stimulation: $U = 17$, p = 0.42, Mann-Whitney *U* test. **b**, Example images of cFos immunofluorescence from stimulated prelimbic cortex tissue in ChR2 mice (right) or eYFP control mice (left). Lower panels depict cells co-labeled for eYFP (green) and cFos (red). Scale = 500 µm (upper), 50 µm (lower). **c**, Example images of cFos immunofluorescence from stimulated animals for statistical analysis in Fig. 9h. LA = lateral, BA = basal amygdala. LHb = lateral habenula. MHb = medial habenula. PVT = paraventricular thalamus. DM = dorsomedial hypothalamus. CPu = caudate putamen. NAcC = nucleus accumbens core. NAcS = nucleus accumbens shell. CA1 = cornu ammonis area 1 of the ventral hippocampus (CA1). DG = dentate gyrus of the dorsal hippocampus. MDL = medial mediodorsal thalamus. MDC = central mediodorsal thalamus. MDL = lateral mediodorsal thalamus. IMD = interomedial dorsal thalamus. Scale = 500 µm (CPu/ NAc images), 200 µm (all other images). Bar graphs depict mean ± SE. Box plots depict median (center line), quartiles, and 10-90% range (whiskers).

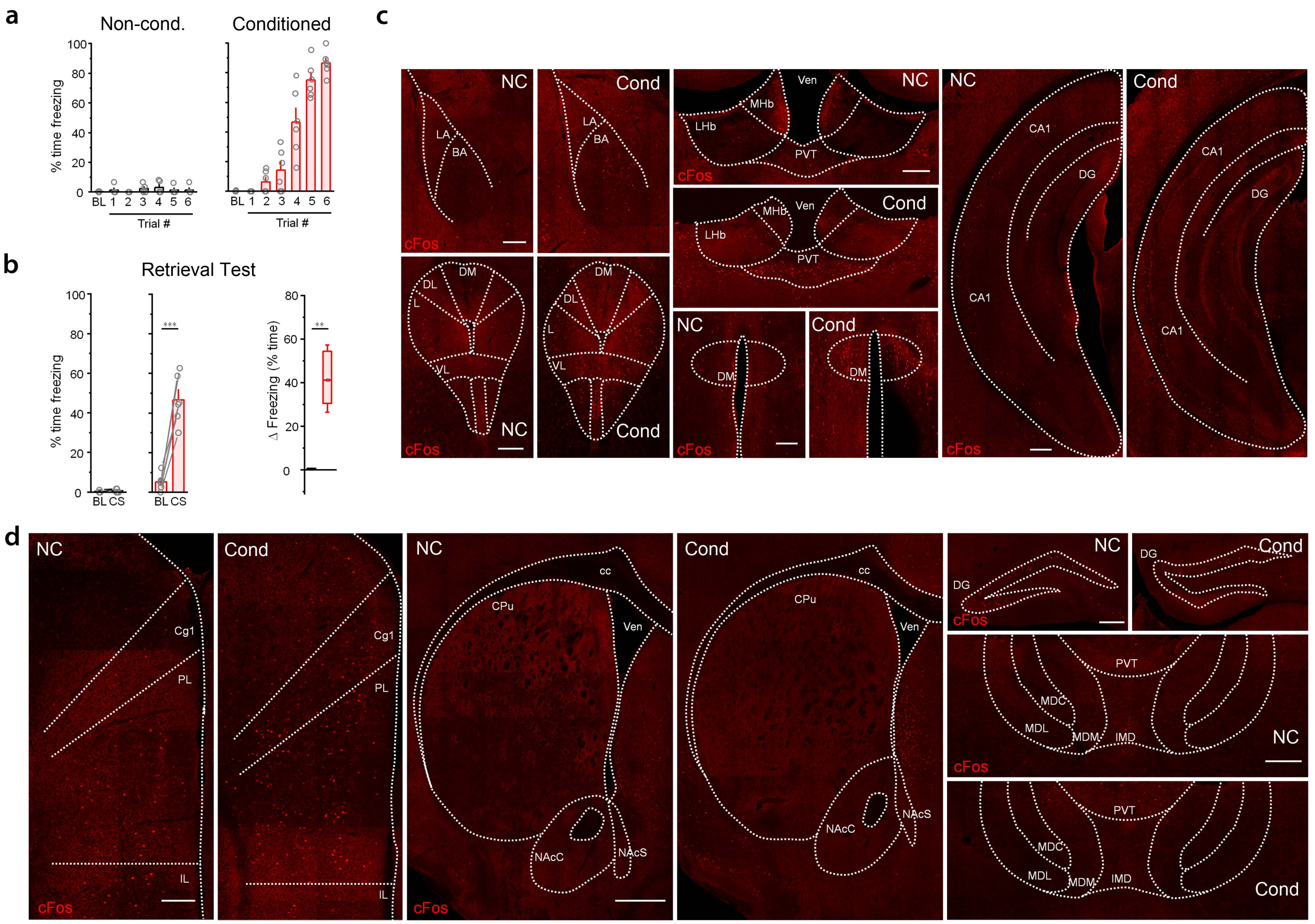


**Supplementary Fig. 16. Behavior and supplementary images for network cFos analysis following memory retrieval.** Data are supplementary to the analysis in Fig. 9i-j. **a**, Percent time freezing during CS-US pairing for conditioned (n = 6) and non-conditioned mice (n = 5). **b**, Percent time freezing during CS retrieval test. Conditioned mice, effect of CS: $t_5 = 8.23$, p = 0.00043, paired t-test. Non-conditioned mice, effect of CS: *W* = 0, p = 0.50, Wilcoxon rank sum test. Conditioned versus non-conditioned mice, change in freezing during CS exposure: *U* = 0, p = 0.0043. **c**, Example images of cFos immunofluorescence following CS exposure in conditioned (Cond) and non-conditioned (NC) mice, for statistical analysis in Fig. 9j. LA = lateral, BA = basal amygdala. LHb = lateral habenula. MHb = medial habenula. PVT = paraventricular thalamus. DM = dorsomedial hypothalamus. CA1 = cornu ammonis area 1 of the ventral hippocampus (CA1). **d**, Additional example images of cFos immunofluorescence following CS exposure in conditioned (Cond) and non-conditioned (NC) mice. CPu = caudate putamen. NAcC = nucleus accumbens core. NAcS = nucleus accumbens shell. DG = dentate gyrus of the dorsal hippocampus. MDM = medial mediodorsal thalamus. MDC = central mediodorsal thalamus. MDL = lateral mediodorsal thalamus. IMD = interomedial dorsal thalamus. Scale = 500 µm (CPu/ NAc images), 200 µm (all other images). **p < 0.01 by paired t-test (**b**, left) or Mann-Whitney *U* test (**b**, right). Bar graphs depict mean ± SE. Box plots depict median (center line), quartiles, and 10-90% range (whiskers).
